# When a chaotropic agent turns into a nutrient – Deciphering the assimilation of guanidine and its utilization to drive synthetic processes in cyanobacteria

**DOI:** 10.1101/2025.03.14.643060

**Authors:** M. Amadeus Itzenhäuser, Andreas M. Enkerlin, Jan A. Dewald, Ron Stauder, Hannes Halpick, Rosalie Schaale, Lisa M. Baumann, Khaled A. Selim, Christina E. Weinberg, Stephan Klähn

## Abstract

Guanidine is a well-known chaotropic agent used to denature proteins and nucleic acids. However, recent studies have demonstrated both the widespread synthesis of guanidine, e.g. in plants and mammals, as well as the widespread occurrence of guanidine metabolism in bacteria, suggesting a broader biological role. Here, we provide insights into guanidine assimilation via guanidine hydrolases (GdmH) in cyanobacteria. The *gdmH* gene is widespread among cyanobacteria and enables growth on guanidine as sole nitrogen source. Strains lacking *gdmH*, naturally or by deletion, failed to grow on guanidine. Expression of *gdmH* increased under nitrogen limitation, regulated by the transcription factor NtcA. However, guanidine is toxic above 5 mM, necessitating GdmH activity and adaptive mutations activating the multidrug efflux system PrqA. The *gdmH* gene is frequently co-localized with ABC transporter genes, which are driven by an additional NtcA-regulated promoter. At low guanidine concentrations, their mutation disrupted guanidine-dependent growth of *Synechocystis* sp. PCC 6803, supporting that they encode a high affinity transport system. In presence of >1 mM guanidine, these mutants grew like wildtype, suggesting the existence of additional uptake mechanisms for guanidine. We next demonstrate the high-affinity binding of guanidine to a previously described, conserved RNA motif located within the *gdmH* 5’ UTR, validating it as a guanidine I riboswitch. By combining it with various promoters, we achieved precise, titratable control of heterologous gene expression in cyanobacteria *in vivo*. Our findings establish guanidine assimilation as an integral element of cyanobacterial nitrogen metabolism and highlight guanidine riboswitches as valuable tools for synthetic biology.

**Significance statement:** Cyanobacteria are promising whole-cell biocatalysts for the sustainable, CO_2_-neutral production of chemicals and fuels. Unlocking this potential requires deep understanding of their metabolism and advanced molecular tools for genetic engineering. We show that cyanobacteria can assimilate guanidine as sole nitrogen source. Because of its toxicity, guanidine metabolism is tightly controlled, involving the transcription factor NtcA and a riboswitch, an RNA element regulating gene expression by guanidine binding. By utilizing this riboswitch, we achieved precise regulation of heterologous genes. Guanidine is inexpensive and effective at low concentrations, making large-scale applications in cyanobacterial cell factories cost-efficient. This study advances our understanding of the metabolic capacities of environmentally important cyanobacteria and their metabolic engineering, highlighting riboswitches as valuable tools for controlling biotechnological processes.

## Introduction

Nitrogen (N) is a central element of life and is incorporated into biomolecules via assimilatory pathways. Although it is highly abundant in Earth’s atmosphere in form of dinitrogen (N_2_), only a few bacteria and archaea are capable of utilizing this chemically inert gas. In contrast, most microorganisms rely on the uptake and assimilation of combined N sources, such as ammonia, nitrate or urea from their environment. Incorporation of ammonium into a carbon skeleton commonly involves ATP-dependent amidation of glutamate, catalysed by glutamine synthetase (GS), yielding glutamine (1, 2). Subsequently, glutamine oxoglutarate aminotransferase (GOGAT) catalyses the transfer of the amide group from glutamine to 2-oxoglutarate, generating two glutamate molecules (3). While the GS-GOGAT cycle is central for ammonium assimilation, additional enzymes enable growth on further N compounds. For example, nitrate is assimilated through a two-step reduction process to ammonium catalysed by nitrate reductase and nitrite reductase (4, 5). Urea can be assimilated via two distinct pathways. Urea either undergoes direct hydrolysis to ammonia and CO_2_, catalysed by urease (6), or it can be carboxylated by urea carboxylase yielding allophanate, which is subsequently hydrolysed to ammonium and CO_2_ by allophanate hydrolase (7).

Another N-rich compound is guanidine, whose biological role has only recently been recognized. Due to its potentially toxic properties, guanidine is usually described to be exported by Gdx-proteins (<u>g</u>uani<u>d</u>ine e<u>x</u>porter proteins) or degraded for detoxification (8, 9). However, there is evidence that guanidine can also be utilized for N assimilation (10, 11), or as sole source of energy and reductant in several bacteria (12). For example, some bacteria can form carboxyguanidine via a guanidine carboxylase. An additional enzyme, a carboxyguanidine deiminase, is then required for the generation of allophanate, which is further metabolised analogously to urea assimilation (9, 13). Moreover, Ni^2+^-dependent guanidine hydrolases (GdmH) capable of directly converting guanidine to urea and ammonia have recently been identified in cyanobacteria (10, 14).

Cyanobacteria are the only prokaryotes performing oxygenic photosynthesis and therefore of critical relevance for the global biogeochemical cycles. In oxygenic photosynthesis, light energy drives the oxidation of water, thereby releasing oxygen as a byproduct (15). The obtained electrons are used to fix and reduce CO_2_ to form carbohydrates and biomass. It is commonly accepted that cyanobacteria caused the rise of free oxygen in the ancient atmosphere, which is known as the great oxygenation event (16–18). Although cyanobacteria are usually associated with photosynthetic CO_2_ fixation and primary production in the oceans, they also contribute massively to the global N cycle by N_2_ fixation or the assimilation of soluble N compounds including nitrate, nitrite, ammonia or urea (19–21). Beyond their environmental impact, cyanobacteria have attracted attention in the pharmaceutical industry as many strains naturally produce active ingredients for medical purposes (22–24). Moreover, cyanobacteria are of increasing biotechnological interest, e.g. as whole-cell biocatalysts for a light-driven, CO_2_-neutral and hence, sustainable production of chemicals and fuel components (25–27).

The utilization of microbes as biocatalysts depends on a comprehensive understanding of their metabolic pathways and regulatory networks to minimize perturbations and to enhance substrate uptake and conversion (28). Accordingly, it is of utmost importance to understand the regulation of major pathways in cyanobacteria including CO_2_ fixation, carbon (C) and N metabolism, which have been investigated for decades. Therefore, a recent report on guanidine-converting enzymes in the cyanobacterium *Synechocystis* sp. PCC 6803 (hereafter *Synechocystis*) and its capability to use guanidine as sole N source (10) was surprising and thus prompts further investigation. In particular, all components for guanidine assimilation need to be identified and integrated into the so-far known regulatory network maintaining the C/N balance. The utilization of various N sources is tightly regulated in bacteria and include various mechanisms targeting the uptake of N compounds and the activity of assimilatory enzymes (1, 2, 29). The control of N assimilation in cyanobacteria includes transcriptional regulation by NtcA. Under N limitation, NtcA binds to a defined DNA motif upstream of various genes encoding, e.g. GS, nitrate reductase as well as transporters for nitrate, ammonium or urea, and activates their transcription (30–32). However, NtcA also represses genes under N limitation, e.g. some encode small proteins that act at the post-translational level to control the activity of other regulatory proteins or enzymes and transporters (33–35). N limitation is perceived intracellularly by the accumulation of 2-oxoglutarate (36) and transduced by a complex signaling cascade involving two major regulatory proteins, PII and PipX (2, 37–39).

To metabolically engineer cyanobacteria, achieving precise control over gene expression and enzyme activities is essential to optimize metabolic flux towards desired products. Riboswitches have emerged as powerful tools in synthetic biology applications to control gene expression also in cyanobacteria (40, 41). Riboswitches are distinct RNA elements mostly found in the 5’-untranslated region (5’UTR) of bacterial messenger RNAs (mRNAs), and work as sensors and signal transducers to control the synthesis of a protein encoded by the same mRNA (42). In particular, secondary RNA structures are re-modulated upon binding of ligands, i.e. metabolites or metal ions, thereby resulting in either activation or inhibition of gene expression (43). The regulation occurs most commonly at the transcriptional level by either forming or resolving an intrinsic terminator or at the translational level by sequestering or releasing the ribosome binding site upon ligand binding (44). Riboswitches tune gene expression according to a metabolic status and thereby play important roles in balancing bacterial metabolism. There are several examples connected to N metabolism, including riboswitches for glycine (45), lysine (46) or glutamine (47), the latter of which are found predominantly in cyanobacteria. Remarkably, a biological role for free guanidine was first noticed by the discovery of different classes of guanidine riboswitches and the subsequent identification of guanidine-converting enzymes in bacteria (9, 48–51).

Here, we have systematically examined cyanobacterial guanidine assimilation. We show that corresponding genes, namely *gdmH* (encoding Ni^2+^-dependent guanidine hydrolase), as well as *hypA2* and *hypB* (encoding maturases for Ni^2+^-incorporation), are widespread in cyanobacteria and allow them to grow on guanidine as sole N source. We show further, that enabling guanidine assimilation by GdmH allows to cope with guanidine toxicity. Coping with guanidine excess also includes its export by the previously characterized multidrug efflux system PrqA that mediates resistance against the herbicide paraquat (methyl viologen) (52, 53). Mutations within the *prqR* (*slr0895*) gene encoding a negative transcriptional regulator for *prqA*, unexpectedly increased guanidine tolerance of the cyanobacterial model *Synechocystis*. Moreover, *gdmH* is frequently associated with genes of an ATP-binding cassette (ABC) transporter, which facilitated the assimilation of guanidine at low concentrations. In *Synechocystis*, *gdmH, hypA2* and *hypB* are transcribed as polycistronic RNA and their expression is induced under N limitation due to NtcA-dependent transcriptional regulation, further supporting a role for guanidine as alternative nitrogen source. Although transcribed from an independent promoter, the ABC transporter genes are similarly controlled by NtcA. Along with that, we experimentally confirmed the presence of an additional transcriptional control of *gdmH* via a guanidine I riboswitch in *Synechocystis* and demonstrate its utilization as a molecular tool to drive heterologous gene expression. By harnessing the regulatory capabilities of guanidine riboswitches, we pave the way for the rational design and optimization of cyanobacterial cell factories for the sustainable production of chemicals and fuels.

## Results

### Genes for guanidine assimilation are widespread among cyanobacteria

The *sll1077* gene (*gdmH*, previously annotated as *speB2*) encodes a Ni^2+^-dependent guanidine hydrolase (GdmH) enabling *Synechocystis* to grow on guanidine as the sole N source (10). To get deeper insight into guanidine assimilation, representative cyanobacteria were searched for the presence of putative *gdmH* homologs. Remarkably, such genes are frequent in cyanobacterial genomes pointing towards a significant biological role of guanidine assimilation in that phylum (**Fig. 1A**). However, many strains also lack *gdmH* genes. At first glance, there is no clear correlation with the different morphological subsections, lifestyles, or physiological traits such as N_2_ fixation. For example, *gdmH* appears to be absent from strains such as *Nostoc* sp. PCC 7120 or *Anabaena cylindrica,* but is present in *Scytonema hofmanni* and *Fischerella thermalis* all of which form heterocysts for N_2_ fixation. Moreover, it is absent from many marine picocyanobacteria of the genus *Prochlorococcus* with their streamlined genomes, but is frequently present in closely related marine *Synechococcus* strains. In contrast, *Synechococcus elongatus* PCC 7942, which is another cyanobacterial model strain, also lacks GdmH.

**Figure 1:**
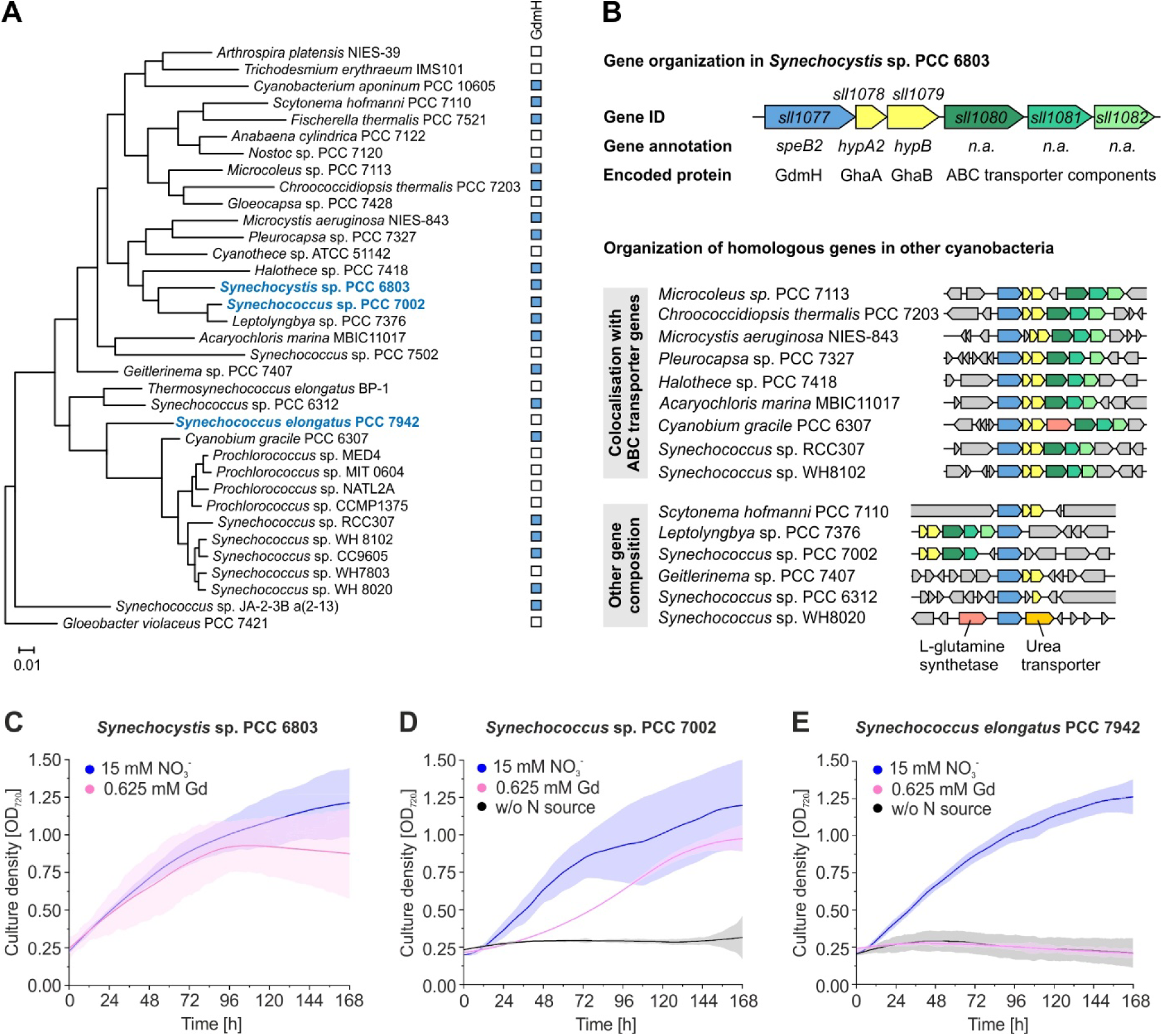
Occurrence of a guanidine hydrolase (GdmH) and associated genes in representative cyanobacteria. **A:** Phylogenetic tree of selected cyanobacteria based on 16S rDNA sequences. Presence of *gdmH* homologs in each strain is shown by a filled square. The strains shown in blue letters were used to analyze guanidine-dependent growth. **B:** Genetic locus encoding GdmH in *Synechocystis* sp. PCC 6803 and organization of homologous genes in other cyanobacteria. The color code of homologs corresponds to the reference genes from *Synechocystis* shown in panel B. **C-E:** Growth of representative cyanobacteria in presence of nitrate (NO_3_^−^) or guanidine (Gd) as sole N source. Growth was automatically monitored in a multicultivator MC-1000 in 10 min intervals. Data are the mean ± SD of triplicates (SD appears as shadow behind mean curves).

Consistent with the Ni^2+^-dependence of GdmH, the corresponding ORF in *Synechocystis* is associated with genes *hypA2* (*sll1078*) and *hypB* (*sll1079*), whose gene products show similarities to the maturase proteins of NiFe hydrogenases responsible for the incorporation of the metal cofactors (**Fig. 1B**). This co-occurrence is common in cyanobacteria although the order of genes can be different from the genetic configuration found in *Synechocystis*. Moreover, the three genes are frequently co-localized with ABC transporter genes with non-specific annotation, i.e. without information about the transported substrate. For example, the gene *sll1080* putatively encodes an aliphatic sulfonate-binding protein of a NitT/TauT family transport system, *sll1081* the corresponding permease and *sll1082* the ATP binding protein. It has been speculated that the *sll1080-sll1082* genes are potentially co-transcribed with *gdmH* and encode a specific ABC transporter for guanidine uptake (10). Their presence has also been noticed in other bacteria such as *Pseudomonas syringae* (12). However, co-transcription, its specificity and its contribution to guanidine assimilation has not been investigated so far (i.e. not experimentally proven yet). Nevertheless, a similar gene organization can be found in many other cyanobacteria harboring a *gdmH* gene, which supports this assumption (**Fig. 1B**). In some cases, *gdmH* is situated in a different genetic environment, but often related to genes for N assimilation. For example, in *Synechococcus* sp. WH8020, *gdmH* is surrounded by genes encoding glutamine synthetase and a urea transporter, further supporting a role in N assimilation.

We aimed at getting further insights into guanidine assimilation and, in particular, guanidine-dependent growth of cyanobacteria. It has previously been demonstrated that *Synechocystis* can grow on guanidine as the sole N source, assimilating it via GdmH (10). Beyond *Synechocystis*, we also tested two other representative model strains, *Synechococcus* sp. PCC 7002 and *Synechococcus elongatus* PCC 7942 (**Fig. 1C-E**). All strains showed similar growth performance under standard conditions, i.e. in presence of sufficient concentrations of nitrate. When nitrate was replaced by guanidine, there was a clear correlation between the presence of *gdmH* and the ability to grow on guanidine. Both *Synechocystis* and *Synechococcus* sp. PCC 7002 harbor a *gdmH* gene and thereby can grow on guanidine (**Fig. 1C, D**). In contrast, *Synechococus elongatus* PCC 7942 lacks a *gdmH* gene and was not able to grow on guanidine, indicating an inability to assimilate it to form biomass (**Fig. 1E**).

### Guanidine toxicity limits its utilization as sole N source

To evaluate the boundaries for guanidine as N source, we monitored growth of *Synechocystis* wildtype (WT) and a *gdmH* knockout strain in the presence of different guanidine concentrations (**Fig. 2**). For the WT best growth performance, i.e. comparable with standard conditions in presence of nitrate, was found at rather low guanidine concentrations between 0.3 and 1.25 mM. However, these low amounts were rapidly consumed as the stationary phase was reached already after 72 hours and the cells subsequently became chlorotic (**Fig. 2C**), which is a clear indication for N limitation in cyanobacteria (54). At 2.5 mM, a greater variation in the optical densities was observed and in presence of 5 mM guanidine no growth could be detected (**Fig. 2C, E**). Nevertheless, it should be noted that cells started growing slowly after a very long lag phase (after a week) pointing towards a compensating mechanism or the emergence of suppressor mutants (see below). In contrast, the Δ*gdmH* mutant did not grow on any of the tested guanidine concentrations (**Fig. 2D, F**).

**Figure 2:**
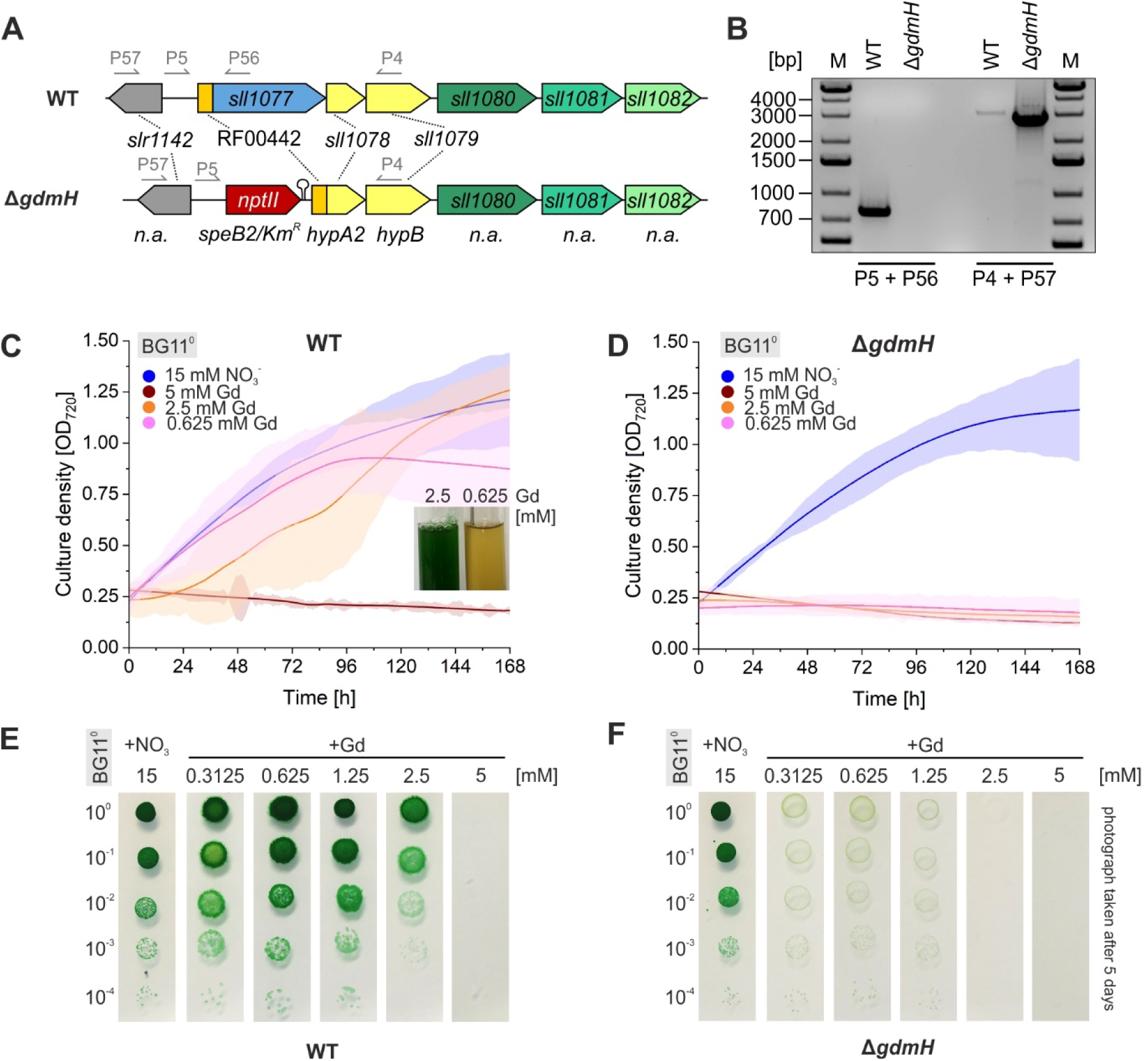
Guanidine-dependent growth of *Synechocystis* sp. PCC 6803. **A:** Genetic configuration of the WT and a *gdmH* deletion mutant (Δ*gdmH*), in which the *gdmH* gene was replaced by a kanamycin resistance cassette. **B**: PCR confirming the corresponding gene deletion in Δ*gdmH*. **C, D:** Growth of WT and Δ*gdmH* in liquid cultures grown in BG11^0^ supplemented with either 15 mM nitrate or different concentrations of guanidine as sole N source. Optical density was automatically monitored in a multi-cultivator MC-1000. Data are the mean ± SD of triplicates. The picture exemplifies culture appearance at the end of the experiment. Yellow pigmentation indicates N limitation induced chlorosis. **E, F:** Photograph of a drop dilution assay of WT and Δ*gdmH* on agar plates supplemented with different N sources and concentrations. Please note that the given N compound represented the sole N source in each case.

Although guanidine can be used by *Synechocystis* as sole N source, it also elicits some toxicity as the WT did not grow at concentrations of 5 mM or above. This was investigated further by experiments in which cells were fed with both nitrate and guanidine (**Fig. 3**). In contrast to our previous observations, growth of *Synechocystis* WT was not or less affected when 5 mM guanidine were added in addition to nitrate that is available in standard BG11 medium (**Fig. 3A, C**). This cannot be explained by a stress avoidance, e.g. by a nitrate-dependent inhibition of guanidine assimilation, as guanidine was still consumed from the medium suggesting its incorporation and processing by an active GdmH (**Fig. 3A**). However, guanidine addition had drastic effects on the Δ*gdmH* mutant that otherwise grew well when nitrate was the sole N source (**Fig. 3B, C**). Even guanidine concentrations as low as 0.1 mM resulted in an immediate growth arrest of the Δ*gdmH* strain. The inability to grow on nitrate when guanidine was also present, underlines its toxicity. Although not as severe as in the Δ*gdmH* mutant, a similar effect was observed in *Synechococcus elongatus* PCC 7942, which naturally lacks the *gdmH* gene. When nitrate-grown cells were supplemented with guanidine, a clear dose-dependent response was observed, with increasing guanidine levels negatively affecting growth (**Fig. 3D**).

**Figure 3:**
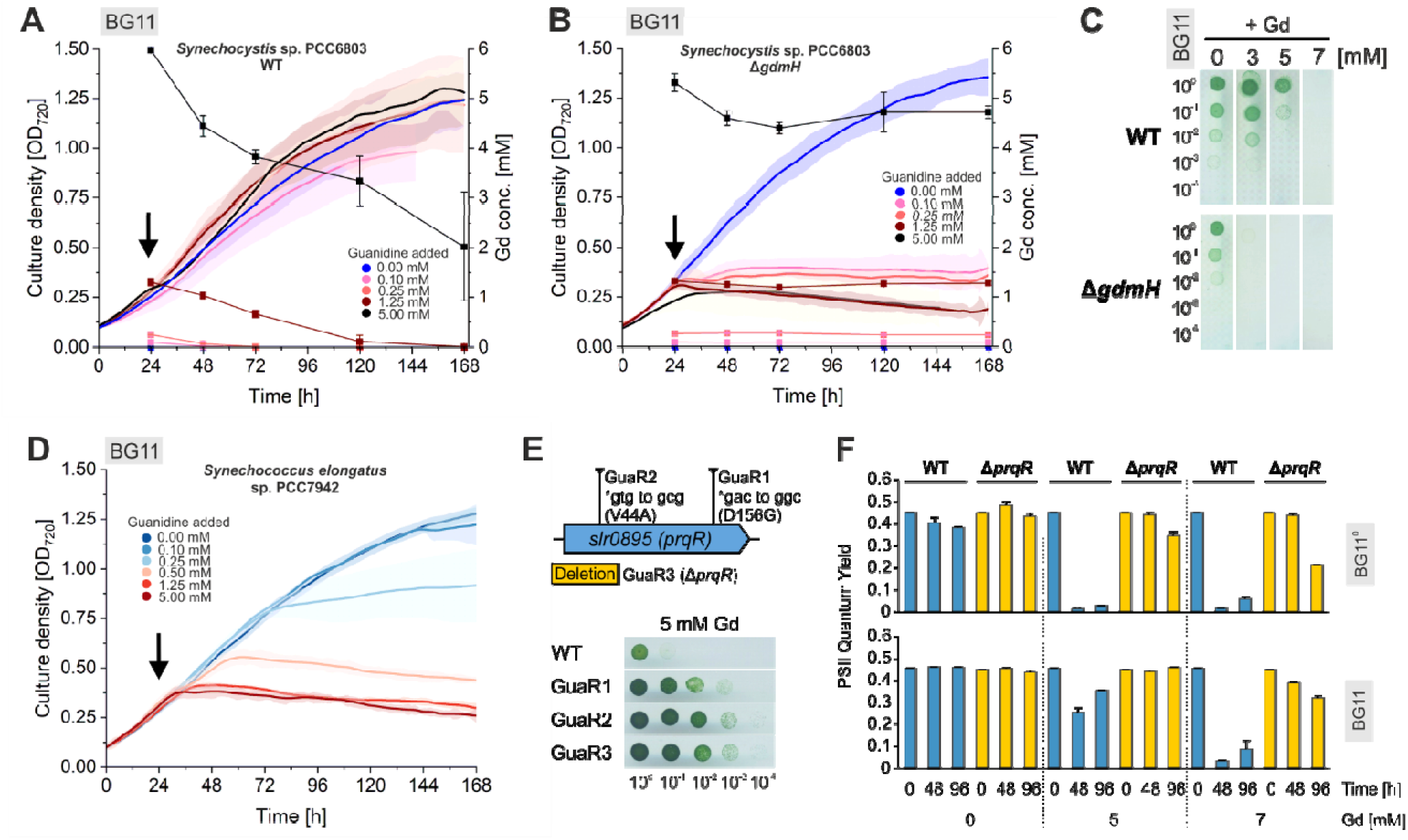
The importance of enzymatic degradation and export in coping with guanidine toxicity. **A-B:** Cultures of *Synechocystis* WT (A) and a *gdmH* deletion mutant (Δ*gdmH*) (B) grown in BG11 medium, i.e. in presence of nitrate to which different guanidine concentrations were added. The arrow indicates the time point of guanidine addition. Growth was automatically monitored as OD_720_ in a multicultivator MC-1000 in 10 min intervals. During the experiments guanidine concentration in the medium was profiled via HPLC and is indicated by rectangles in the respective color of initial guanidine concentration. **C:** Drop dilution assay of *Synechocystis* WT and Δ*gdmH* on BG11 agar, i.e. in presence of nitrate, supplemented with different guanidine concentrations. **D:** Growth of *Synechococcus elongatus* PCC 7942 in BG11 supplemented with additional guanidine (same conditions as described for panel A & B). **E:** Genetic locus of the *prqR* gene in which various mutations have been identified by sequencing the genomes of GuaR1-3 and drop dilution assay of suppressor mutants GuaR1-3 grown on BG11 agar in presence or absence of 5 mM guanidine. **F:** Quantum yield of photosystem II in cells of *Synechocystis* WT and GuaR3 (Δ*prqR*) that have been incubated in BG11 or BG11^0^ in presence of different guanidine concentrations for 48 and 96 h, respectively. Data are the mean ± SD of triplicates.

Guanidine toxicity indicates that GdmH cannot sufficiently degrade it at elevated concentrations. However, on agar plates resistant colonies emerged when *Synechocystis* WT was incubated with 5 mM guanidine for a prolonged period of time. We picked three resistant colonies and sequenced their genome. In all three resistant strains (GuaR1-3), the only mutated gene was *slr0895,* annotated as *prqR* (**Fig. 3E**). While strains GuaR1 and GuaR2 harbored amino acid substitutions (D156G and V44A, respectively), strain GuaR3 showed a 199 bp deletion including the proximal 165 bp of the *slr0895* ORF. The corresponding gene product PrqR is a transcriptional repressor of the multidrug efflux transporter PrqA encoded downstream by *slr0896* (55, 56). The latter was reported to be upregulated in *prqR* mutants (53, 56). Therefore, *prqA* gene expression is likely increased in GuaR1-3, in turn leading to detoxification of the cells through removal of excess intracellular guanidine. The three (GuaR1-3) strains were highly resistant to guanidine compared to WT cells (**Fig. 3E**). We further characterized the deletion variant GuaR3 (Δ*prqR*) using PAM fluorometry and calculated the photosystem II (PSII) quantum yield as a measure for photosynthetic performance after cells were incubated with guanidine. Compared to WT, the Δ*prqR* mutant indeed showed an increased tolerance to higher concentrations of guanidine, regardless of nitrate presence (**Fig. 3F**). Even after 4 days in presence of 7 mM guanidine, the Δ*prqR* mutant was able to perform photosynthesis, whereas WT cells were strongly impaired at 5 mM, especially in absence of nitrate in accordance with growth (**Figs. 2E and 3A**). Expulsion of excess guanidine by dysregulated PrqA probably alleviates the workload of GdmH, allowing it to hydrolyze the remaining guanidine to be used as an N source.

*Guanidine uptake via an ABC transporter precedes its assimilation by enzymatic degradation* Next, we analyzed the impact of disrupting the associated ABC transporter genes on guanidine-dependent growth of *Synechocystis*. We focused on the two genes *sll1080* and *sll1081*, which encode a potential substrate binding protein for guanidine (*sll1080*) and a permease that might transport guanidine through the membrane (*sll1081*). Both genes were disrupted by gene cassettes encoding resistance against kanamycin or spectinomycin (**Fig. 4A**). The mutants were obtained via homologous recombination and complete replacement of the native gene loci was verified by PCR (**Fig. 4B**). At a first glance, considering colony formation on agar plates, strain Δ*sll1080* that lacks the substrate binding protein showed no obvious growth limitation when guanidine was the sole N source (**Fig. 4C**). However, in presence of low guanidine concentrations (< 0.5 mM) the initially green cell material turned yellowish after a couple of days, which was not the case for the WT. As a chlorotic phenotype indicates N limitation, these observations suggest that strain Δ*sll1080* is at least partially affected in assimilating guanidine compared to WT, presumably due to limited guanidine uptake. Nevertheless, this phenotype could be compensated by guanidine concentrations of >1 mM as both WT and Δ*sll1080* cells stayed green (**Fig. 4C**). Growth performance and guanidine assimilation was further analyzed using liquid cultures. Under standard growth conditions, i.e. in BG11 medium containing 17.5 mM nitrate, both mutants Δ*sll1080* and Δ*sll1081* grew similar to the WT confirming that they are not generally essential (**Fig. 4D**). Interestingly, both mutants showed faster growth in presence of 2 mM guanidine as sole N source, consistent with a faster consumption of guanidine from the medium (**Fig. 4E**). However, this became inverted when guanidine was used in limiting concentrations. In presence of 0.5 mM guanidine, the WT reached the maximum OD between 48 and 70 hours after inoculation and then became chlorotic due to N limitation as guanidine was entirely consumed. In contrast, both mutants grew significantly slower, consistent with still detectable amounts of guanidine in the medium after a week (**Fig. 4F**). In presence of 0.1 mM guanidine, both Δ*sll1080* and Δ*sll1081* were not able to grow and did not deplete guanidine from the medium, while the WT cells consumed all guanidine within 24 h (**Fig. 4G**). Based on these data, we concluded that the respective ABC transport system is crucial when only trace amounts of guanidine are available. The Sll1080 substrate binding protein presumably has a high affinity to guanidine, similar to other SBPs of various ABC transporters (57, 58). Nevertheless, there seems to be another guanidine uptake mechanism that sufficiently compensates *sll1080/1081* mutations in presence of higher external concentrations (i.e. a low affinity system). Faster growth of Δ*sll1080* and Δ*sll1081* at 2 mM guanidine may also indicate an internal regulatory crosstalk that aims at compensating the lacking ATP-dependent guanidine import.

**Figure 4:**
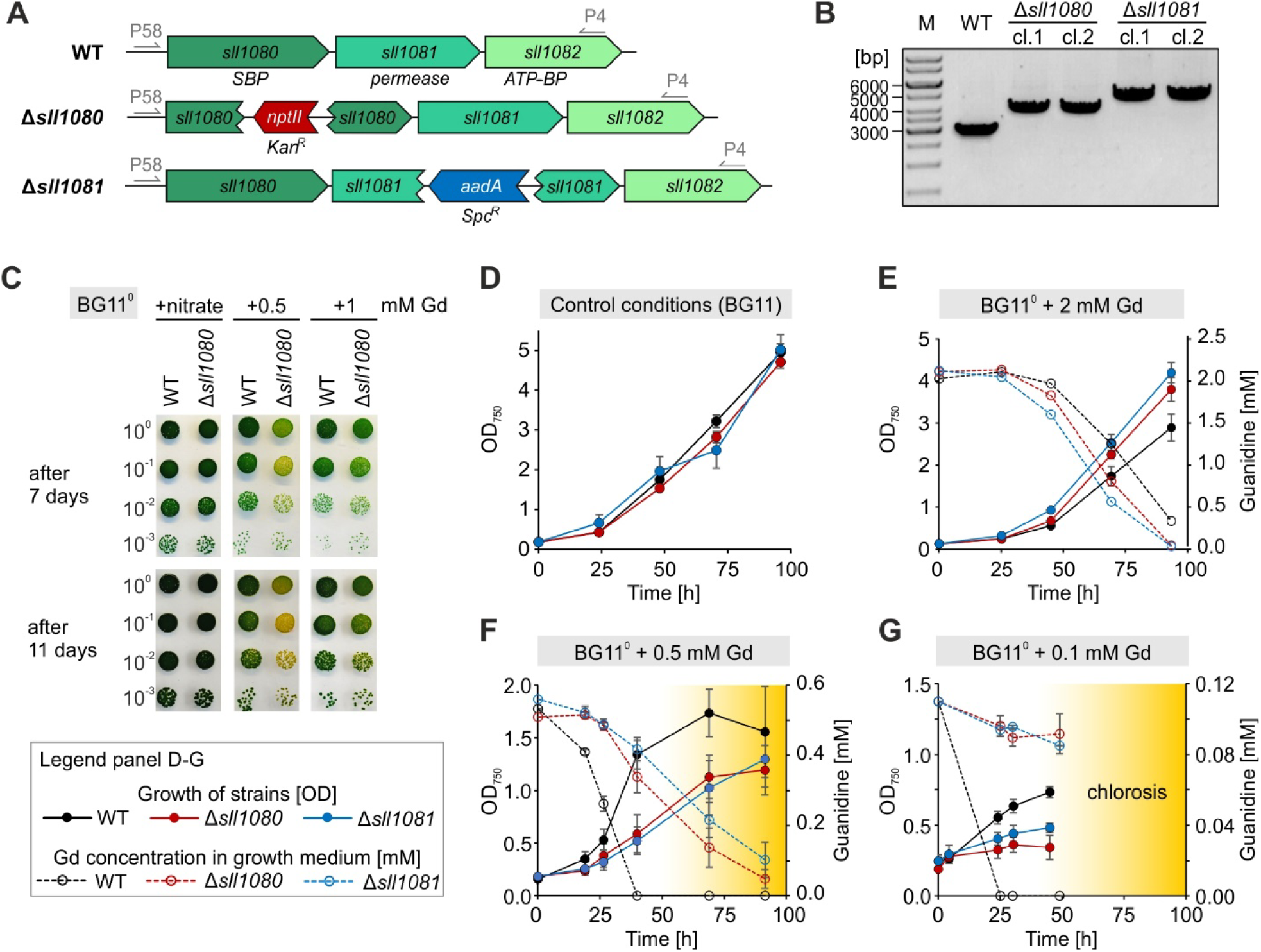
Functional analysis of ABC transporter genes associated with *gdmH* in *Synechocystis*. **A:** Genetic configuration in the WT and two mutants, in which the ORFs *sll1080* or *sll1081* have been interrupted by antibiotic resistance cassettes. The arrows indicate primer binding sites used to verify mutagenesis and full segregation in the obtained clones. **B:** Agarose gel analysis of PCR products confirming disruption of the targeted genes in two independent clones (cl.). As *Synechocystis* is polyploid, full segregation of the mutant alleles was also confirmed by these data as no PCR product corresponding to the WT allele was detected in Δ*sll1080* and Δ*sll1081* mutant cells. **C:** Drop dilution assay for the WT and Δ*sll1080* on agar plates containing BG11^0^ supplemented with nitrate (control) or 0.5 and 1 mM guanidine (Gd), respectively. Photographs were taken after 7 and 11 days. Yellowish appearance of cells indicates chlorosis induced by N limitation. **D-G:** Growth performance of mutants Δ*sll1080* and Δ*sll1081* compared to the WT. Growth was monitored as OD_750_ (solid lines, left y axis) in Erlenmeyer flask cultures, either using standard BG11 medium containing nitrate or in BG11^0^ supplemented with guanidine as sole N source, as indicated. Guanidine consumption from the medium has been monitored by HPLC (dotted lines, right y axis). Please note that for 0.1 mM guanidine growth monitoring has only been conducted until 50 h as the cells already started bleaching due to N limitation. Data are the mean ± SD of three biological replicates.

### Transcriptional organization and regulation of genes for guanidine assimilation

Data from previous transcriptome analysis (59), allowed a closer investigation of the potential transcriptional units and promoter organization of *gdmH* and associated genes (**Fig. 5A**). It has been postulated that *gdmH* forms an operon with the downstream genes encoding the maturases for Ni^2+^-incorporation as well as the ABC transporter system (10). Indeed, a polycistronic transcript covering *gdmH* (*sll1077*), *hypA2* (*sll1078*) and *hypB* (*sll1079*) has been mapped and corresponds to transcriptional unit (TU) 795 (**Fig. 5A**). However, the genes *sll1080-sll1082* appear to form another operon that is transcribed independently from TU795 (59). This is consistent with another transcriptional start site (TSS) present upstream of *sll1080* giving rise to the independent TU794 covering *sll1080-sll1082* (**Fig. 5A**). Similar to other genes involved in N assimilation such as *glnA* that encodes glutamine synthetase, all genes showed clear upregulation under N depletion (**Fig. 5B**). However, the pattern of upregulation was distinct for both TUs, supporting the assumption of independent transcription and regulation of *sll1077-sll1079* and *sll1080-sll1082* (**Fig. 5B**). An independent transcriptional organization and upregulation under N limitation is also consistent with the presence of a DNA motif that partially matches the consensus binding sequence of NtcA (GTA-N_8_-TAC, (30)) upstream of both *sll1077* and *sll1080*. Moreover, these putative NtcA binding sites center around position −42 (referring to TSS, +1; **Fig. 5C**). This promoter organization resembles the situation upstream of other N-responsive genes including *glnA*, whose transcription is activated by NtcA when N is limiting (31, 60). Consistently, ChIPSeq analysis confirmed that NtcA binds upstream of *gdmH* (*sll1077*) under N limitation (32). As this has not yet been reported for the motif upstream of *sll1080*, we performed a promoter test assay to verify the presence of a controllable promoter as well as NtcA-dependent activation of gene expression under N limitation. For this, we fused the upstream sequence of *sll1080* with a *superfolder gfp* (*sfgfp*) gene and monitored the fluorescence of the corresponding gene product *in vivo*. We also included a promoter variant in which the putative NtcA binding motif was point mutated (**Fig. 5C**). The genetic configuration of the generated reporter strains is given in **Fig. 5D**. Indeed, strain P80 showed significant GFP fluorescence compared to a negative control strain that did not harbor a *sfgfp* gene. Consistent with NtcA-dependent regulation, the GFP signal increased by a factor of 10 under N limitation, which was not the case for strain P80M carrying the mutated promoter variant (**Fig. 5E**). These data confirm the presence of an active promoter upstream of *sll1080*, and further support that the *sll1080-sll1082* genes are indeed transcribed independently from *gdmH* and are nonetheless controlled by NtcA.

**Figure 5:**
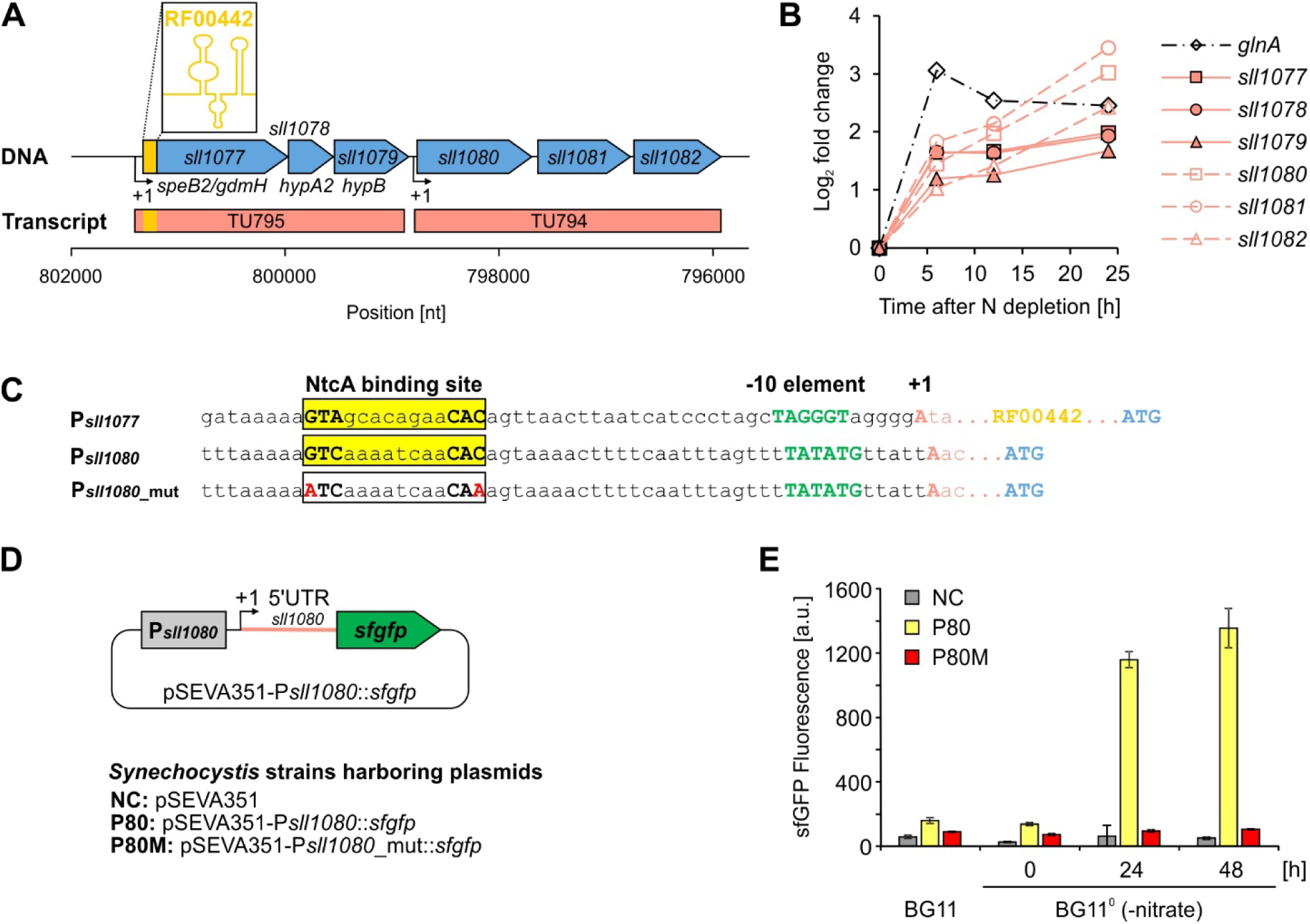
Transcriptional organization of genes for guanidine assimilation in the cyanobacterium *Synechocystis* sp. PCC 6803. **A:** Genetic locus encoding guanidine hydrolase (GdmH) and associated proteins. The *gdmH* gene is transcribed as a polycistronic RNA (transcriptional unit 795; TU795) together with genes *hypA2* and *hypB* encoding maturase proteins for the incorporation of Ni^2+^ co-factors required for guanidine hydrolysis. Moreover, TU795 is associated with an RNA motif (RF00442) predicted to function as the aptamer of a guanidine riboswitch. Downstream of *gdmH, hypA2 and hypB*, another TU is transcribed (TU794) covering genes *sll1080-sll1082*, encoding a guanidine ABC transporter. **B:** Gene expression profile for *sll1077*-*sll1082* genes in response to N limitation (-N). Data for each gene have been extracted from previous transcriptomes of N-depleted *Synechocystis* cells (93). **C:** Promoter sequences found upstream of the genomic region covered by TU795 and TU794, respectively. A common - 10 element as well as potential binding sites for the major transcription factor of N regulated genes, NtcA, are present. Data for the transcriptional start sites (TSS, +1) and promoter annotation were extracted from literature (59, 94). **D:** Genetic configuration of reporter strains harboring pSEVA351 plasmid derivatives with a *sfgfp* gene fused to the upstream sequence of *sll1080* (P*sll1080*) or a mutated variant (P*sll1080*_mut), resulting in the strains P80 and P80M, respectively. A strain lacking *sfgfp* served as negative control (NC). **E:** GFP fluorescence in cells of reporter strains that were cultivated under standard growth conditions (BG11, = 17.5 mM nitrate) or incubated in nitrate-depleted medium (BG11^0^) for 48 h. Data are the mean ± SD of three biological replicates with three technical replicates of each.

In addition to transcriptional regulation by NtcA, a conserved RNA motif (RF00442) is part of TU795 and situated directly upstream of the *gdmH* (*sll1077*) open reading frame (**Fig. 5A**). According to the RFAM database, RF00442 represents the aptamer domain of a guanidine I riboswitch that controls gene expression in a guanidine-dependent manner (9). However, no experimental evidence for this regulation has been provided for cyanobacteria yet, particularly for *Synechocystis*. To validate the interaction between guanidine and the conserved RNA, in-line probing (ILP) experiments were performed (**Fig. 6**). ILP exploits the fact that regions of the RNA that are unpaired and thus more flexible are more prone to self-cleavage initiated by the 2[-hydroxyl group of the RNA. Therefore, ILP can be used to gain information about the RNA structure and possible ligand interactions (61). ILP of the WT RNA showed that the aptamer domain folds into the secondary structure consensus of guanidine I riboswitches and revealed structural modulation of regions A54 to A57 and A98 to G100 with increasing guanidine concentrations (**Fig. 6A, B**). These findings strongly suggest an interaction between the RNA and guanidine and are in accordance with data of the guanidine I riboswitch of *Nitrosomonas europaea*, where modulation was observed in the same regions of the RNA (9). To prove the specificity of these interactions, the RNA variant M1 was tested (**Fig 6A**), which bears a mutation at a highly conserved position within the RNA aptamer that was previously shown to interrupt ligand binding in guanidine I riboswitches (9). ILP analysis of variant M1 showed no modulated regions in the presence of guanidine indicating a lack of binding (**Fig. S1**). Quantitative analysis of the band intensities of the riboswitch RNA enables the calculation of the dissociation constant (K_d_) for the interaction with guanidine resulting in a K_d_ of ~58 µM for the WT variant (**Fig. 6C**), in accordance with the affinity of the previously studied guanidine riboswitch of *N. europaea* (60 µM) (9). In contrast, quantification of the cleavage signals of the corresponding regions in the M1 mutant further underlined that RF00442-M1 is not able to bind guanidine compared to WT (**Fig. S1**). Thus, our ILP results confirmed the interaction of guanidine with RF00442 found upstream of *gdmH*.

**Figure 6:**
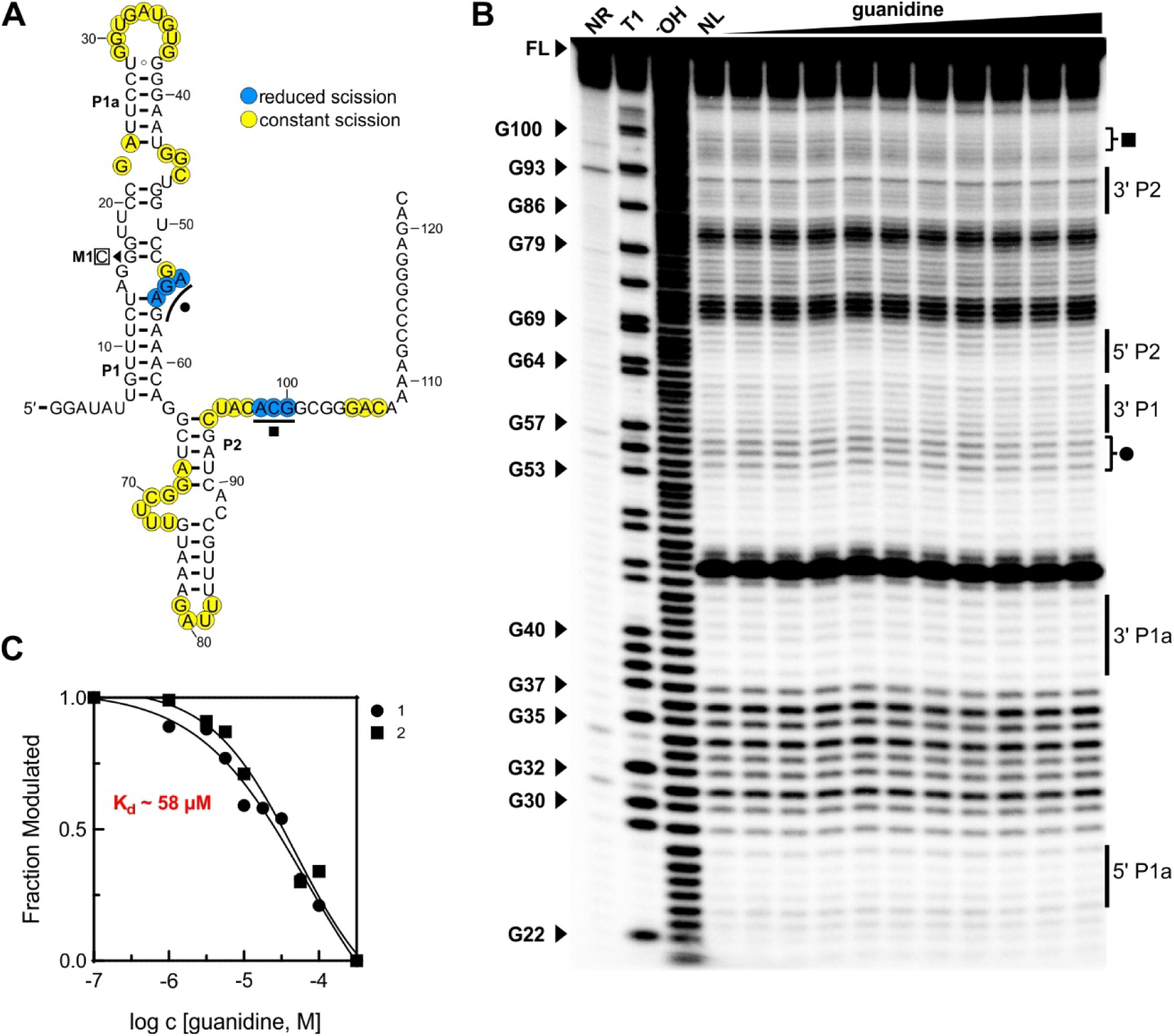
Guanidine binding by the predicted aptamer in the untranslated region of the *gdmH* transcript. **A:** Sequence and secondary structure of RF00442-WT. Colored nucleotides represent ILP cleavage pattern. Modulated regions during in-line probing were marked as in panel B. **B:** In-line probing (ILP) analysis of RF00442-WT. Guanidine concentrations from 100 nM to 316 µM were tested. NR – no reaction control, T1 – RNase T1 digestion, ^−^OH – partial alkaline hydrolysis, NL – no ligand control. Regions with guanidine-dependent conformational changes are marked (Circle and Square). **C:** Graph of the K_d_ analysis of RF00442-WT using pixel analysis of band regions labeled with the same symbols as in panel B.

### Making use of a guanidine riboswitch to enable cheap induction of gene expression for biotechnological applications in cyanobacteria

To test if the RF00442 motif enables guanidine-dependent gene expression in *Synechocystis,* we fused it to a *sfgfp* gene and the promoter P_J23101_ that is considered to be constitutively active in *Synechocystis* (62). The latter has been used in a similar way to analyze the function of glutamine riboswitches (47). In particular, we made use of the 5’ UTR of *gdmH* (*sll1077*) that harbors RF00442 as well as the regulatory sequences for translation initiation. The P_J23101_::RF00442::*sfgfp* construct was introduced into plasmid pSEVA351, which replicates in *Synechocystis*. We generated three *Synechocystis* strains that harbor different plasmid derivatives and/or RF00442 variants: a control strain (CS) harboring pSEVA351 with the described construct and a non-modified RF00442 motif, a mutant strain M1 harboring the same plasmid but with a point mutated RF00442(G16C) that cannot bind guanidine (**Fig. 6A**) and a negative control (NC) strain only harboring a non-modified pSEVA351 plasmid, i.e. without a *sfgfp* gene (**Fig. 7A**). The generated reporter strains were analyzed for GFP fluorescence in the presence or absence of guanidine. Indeed, a specific guanidine-responsive fluorescence signal was obtained via fluorescence microscopy that was absent in cells of the NC strain lacking the P_J23101_::RF00442::*sfgfp* construct (**Fig. 7B**). Next, we quantified the GFP signals using fluorescence spectroscopy (**Fig. 7C**). Cells were grown in BG11, i.e. in presence of 17 mM nitrate and were then additionally exposed to 5 mM guanidine. Although significant GFP fluorescence was obtained in the cells of CS under control conditions compared to the NC strain, the signal increased by a factor of 4-5 upon addition of guanidine (applied as guanidine HCl, **Fig. 7C**). This did not occur when the same concentration of HCl was added, excluding that the GFP signal was induced by a chloride response. Moreover, in strain M1 harboring a mutated RF00442 motif that does not bind guanidine (see **Fig. 6**), no increase in GFP fluorescence could be detected (**Fig. 7C**). This observation confirmed that guanidine exclusively stimulates *sfgfp* gene expression, mediated by a guanidine riboswitch. To analyze the dynamic range of riboswitch-mediated gene expression, we performed a dose-response analysis. For this purpose, we pre-cultivated cells of the CS in BG11 and treated it with different guanidine concentrations. Already at a concentration of 100 µM a clear induction of *sfgfp* gene expression was detectable (**Fig. 7D**). A saturation was reached at 750 µM and a clear drop to 50 % of the maximal fluorescence was obtained for 5 mM guanidine. This is consistent with the observed guanidine toxicity at high guanidine concentrations (see **Fig. 2**). Collectively, we concluded that a functional guanidine riboswitch is present upstream of *gdmH,* which enables guanidine-specific gene expression. This guanidine riboswitch is rather responding to low guanidine concentrations, further supporting the notion of evolving an assimilation system composed of a guanidine degrading enzyme along with a high affinity ABC-guanidine transporter to enable its utilization as an alternative N source under nitrogen limiting conditions. The fast response of the guanidine riboswitch is also attractive for biotechnological applications.

**Figure 7:**
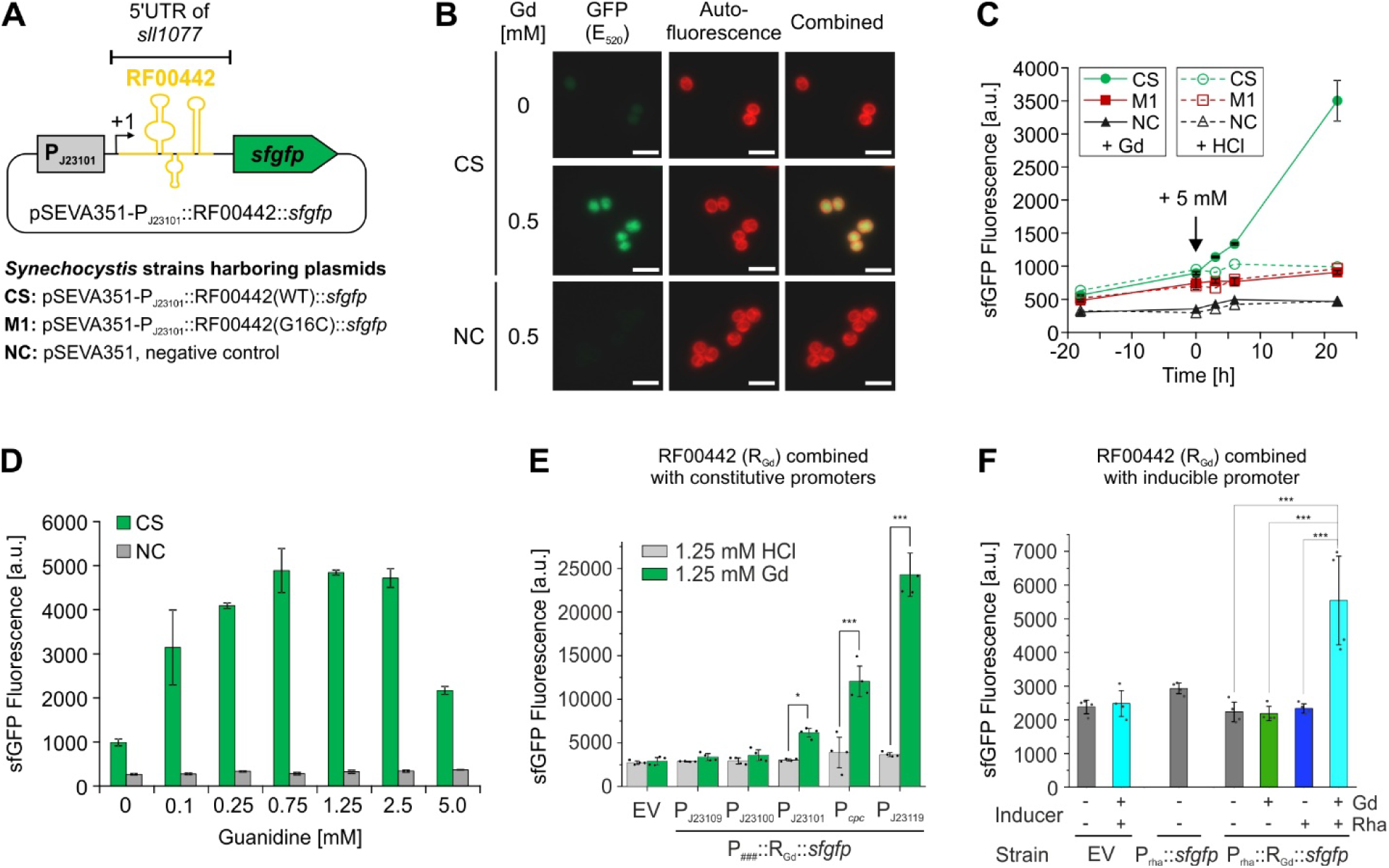
Guanidine-dependent gene expression mediated by a guanidine I riboswitch in *Synechocystis*. **A:** Design of reporter strains harboring derivatives of the replicative plasmid pSEVA351 enabling transcription of a *sfgfp* gene driven by the strong constitutive promoter P_J23101_. In between, the 5’UTR of the gene *sll1077* was introduced that harbors the RNA motif RF00442 resembling a guanidine-binding aptamer. In addition to the control strain (CS) with a WT variant of RF00442, a mutant strain (M1) was generated in which G at position 16 was replaced by C (mutation M1 is the same as used for in-line probing). A strain harboring a pSEVA351 plasmid lacking the reporter construct served as negative control (NC). **B:** Microscopic pictures of recombinant *Synechocystis* cells that were grown in presence or absence of guanidine (Gd), which was applied as Gd hydrochloride. The scale bar corresponds to 5 µm. **C:** Kinetics of GFP fluorescence *in vivo* in response to guanidine. The arrow indicates the time point when, Gd or HCl were added to the cultures at a final concentration of 5 mM. **D:** GFP fluorescence of the control strain that was grown in presence of different Gd concentrations for 24 hours. **E:** Fluorescence signals obtained for *Synechocystis* cells harboring different pSEVA351-based plasmid derivatives in which an *sfgfp* gene was fused to the guanidine riboswitch whereby transcription was driven by constitutive promoters of different strength. Reporter signals were compared for cells supplemented with 1.25 mM guanidine (induction) or HCl (no induction; negative control) for 24 hours. **F:** Fluorescence signals obtained for *Synechocystis* cells that harbor a SEVA351 plasmid with a *sfgfp* and in which the guanine-controllable riboswitch was combined with promoter P*rha* that enables rhamnose-inducible transcription. Corresponding inducer presence (+) or absence (-) is indicated. Data are the mean ± SD of three biological replicates with three technical replicates of each.

Guanidine is a rather cheap chemical and hence could be used as an inducer to control processes based on microbial cell factories at large scale without significantly raising the cost of a potential production process. The guanidine I riboswitch could be an effective molecular tool as it senses and responds to quite low concentrations and allows titration of gene expression. Here, we provide a series of replicative plasmids that harbor different promoters combined with the Gd riboswitch (**Fig. S2A**). These plasmids facilitate the expression of genes in a guanidine-dependent manner with also considering different strengths of expression and a tight control to avoid leaky expression. The plasmid series is based on pSOMA17 and pSEVA351, which are compatible with each other and can therefore be maintained together, in a single cell line (63), increasing the potential for combinatorial engineering. The plasmids harbor a multi cloning site for the simple introduction of genes equipped with sequences allowing the straight forward addition of a His- or FLAG tag for immunodetection (**Fig. S2B,C**).

Similar to the observations made for the P_J23101_::RF00442::*sfgfp* construct, guanidine-dependent GFP fluorescence was detected in strains harboring fusions of RF00442 with other constitutive promoters (**Fig. 7E**). P_J23101_ turned out to be of intermediate activity compared to P_cpc_ or P_J23119_, which resulted in a 2-fold and 5-fold higher GFP fluorescence (**Fig. 7E**). In contrast, the strains harboring fusions of RF00442 with P_J23109_ and P_J23100_ showed rather low GFP fluorescence. But in each case, an increase was observed upon guanidine addition even though the promoters are considered to be constitutively active (**Fig. 7E**). Nevertheless, low GFP fluorescence was detected under control conditions without guanidine that was absent in cells of negative control strains lacking a *sfgfp* gene construct. The riboswitch alone is therefore somewhat leaky, which is problematic in a process that needs to be tightly controlled. Therefore, we also combined it with P_rha_, a controllable promoter that responds to rhamnose (64). This configuration appears to be entirely OFF under control conditions and can be activated by adding rhamnose and guanidine (**Fig. 7F**).

The described plasmid series enables guanidine inducible gene expression and opens up a new possibility of managing a bioprocess. As guanidine can induce gene expression in different levels (**Fig. 7E**) and simultaneously can serve as the sole N source for growth (**Fig. 2C**), a switch from nitrate to guanidine in the medium can be used to control the transition from a growth to a production phase. Exemplarily, we simulated a steady state process in which BG11 medium (nitrate as N-source) was used to grow the cells to the targeted biomass density, then followed by continuous feeding of nitrate-free BG11^0^ medium supplemented 5 mM guanidine to keep the biomass density stable (see **Fig 8A** for a schematic representation of the process). The reached guanidine concentration resulting from the feed allows cell maintenance and simultaneously induces heterologous gene expression. The latter encodes for an enzyme, which e.g. allows the transformation of a co-fed substrate to a desired product (**Fig 8B**). By coordinating the guanidine concentration in the feed medium and the targeted biomass density, the resulting steady state concentration of guanidine for induction of gene expression can be adjusted.

**Figure 8:**
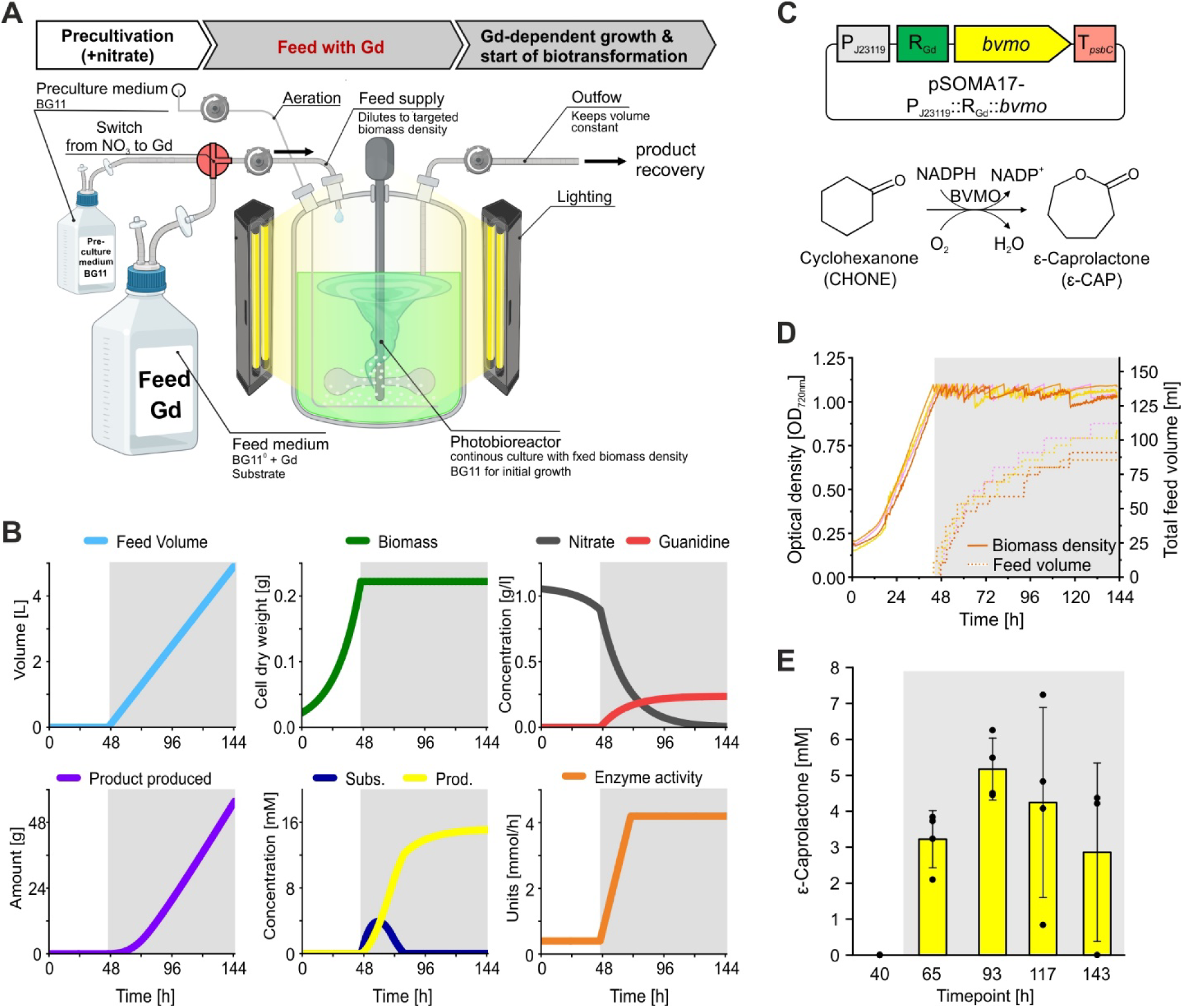
The dual role of guanidine as N source and inducer of gene expression allows a new management of bioprocesses with cyanobacteria. **A:** Schematic representation of a bioreactor in a steady state process with a constant biomass. While the cells are grown to the targeted biomass density in BG11 medium with nitrate as the N-source, the feed medium to constantly dilute the culture to the targeted biomass density contains guanidine as N-source. The change of the N source from nitrate to guanidine can be used to simultaneously induce expression of target genes. Parts of this panel were generated using BioRender. **B:** Computational simulation of selected process parameters in 1 L culture volume with an OD_750_ of 1, 17 mM nitrate in the initial medium, 5 mM guanidine and 15 mM substrate in the feed medium, and 70 U maximal specific activity for the biotransformation. The grey shading illustrates the phase of the guanidine feed. The simulation was conducted using Berkley Madonna. **C:** Scheme of a plasmid from the generated plasmid library that was equipped with a gene for a Baeyer-Villiger monooxygenase, enabling the biotransformation of cyclohexanone to ε-caprolactone. A *Synechocystis* strain carrying this plasmid was used for the *in vivo* demonstration of the simulated process. **D:** Biomass density monitored as optical density at 720 nm and feed volume over the time of the bioprocess for four independent cultures. **E:** Concentration of ε-caprolactone at different timepoints in a bioreactor which was managed in a steady state mode with guanidine in the feed medium as simulated. Data are the mean ± SD of four independent cultures.

The feasibility of such a process design was demonstrated using a *Synechocystis* strain harboring a plasmid coding for a Baeyer-Villiger monoxygenase (BVMO), a notable biocatalyst transforming cyclohexanone to ε-caprolactone (65, 66). The plasmid was generated from the library presented above and was designed in a way that the corresponding *bvmo* gene was controlled by the P_J23119_:: RF00442 fusion promoter (**Fig. 8C**). In a steady state process as described before, growth of the strain could be maintained after the switch of the N-source from nitrate to guanidine (**Fig. 8D**) while ε-caprolactone, the desired product of the biotransformation, could be produced continuously from cyclohexanone which was co-fed (**Fig. 8E**). Thereby, a concentration of guanidine feasible for gene induction was reached in the medium. In contrast, in an analogous process with nitrate instead of guanidine in the feed medium, no notable amount of guanidine for induction of enzyme expression was detected resulting in a drastically lower product concentration over the complete process time (**Fig. S3**).

## Discussion

### The biological role of guanidine

Guanidine is commonly used as chaotropic agent to denature macromolecules such as proteins. Due to the strong interaction of guanidine with a variety of amino acids, it also shows inhibitory effects on enzymes, particularly with acidic amino acids in their active site even at low concentrations (67). Nevertheless, guanidine is part of several biomolecules. Chemicals containing guanidine moieties are commonly used as drugs in medicine, due to their antimicrobial and antifungal properties (68). Even though microbial degradation of guanidine was observed in a variety of surface water samples already in the 1980s (69), a biological role of free guanidine remained ambiguous until guanidine-specific riboswitches were revealed in various bacterial species (9, 48–51). A conserved guanidine-sensing mechanism pointed towards a significant biological role, consistent with the subsequent discovery of guanidine-degrading enzymes. As reported here, a functional guanidine I riboswitch is located upstream of a gene encoding a Ni^2+^-dependent guanidine hydrolase (GdmH) in cyanobacteria, exemplified for the model strain *Synechocystis*. The *gdmH* gene is part of the NtcA regulon and is up-regulated under N limitation as many other genes involved in N assimilation (32). However, it requires guanidine to obtain large amounts of a functional GdmH enzyme due to the fact that even high rates of transcription initiation by strong promoters only resulted in basal expression as long as guanidine was absent. This differential regulation is achieved through the action of the guanidine I riboswitch that turns on *gdmH* expression in the presence of guanidine. Previously studied guanidine I riboswitches that contain the same highly conserved nucleotides function by disrupting an intrinsic transcription terminator stem upon guanidine binding (9, 70, 71). For example, the guanidine I riboswitch from *Desulfotomaculum ruminis* showed in single-round transcription termination assays significantly less terminated transcription product when guanidine was present (9). Furthermore, X-ray crystallography data of guanidine I riboswitches from *Sulfobacillus acidophilus* (70) and *Dickeya dadantii* (71) show evidence for a terminator stem disruption in the ligand-bound state. Therefore, the high sequence and structural similarity to guanidine I riboswitches of other bacteria suggests that the riboswitch from *Synechocystis* regulates gene expression at the transcriptional level as an “ON”-switch rather than at the translational level.

The capability of degrading guanidine to urea and ammonium allows *Synechocystis* and other cyanobacteria to grow on guanidine as sole N source. In nature, guanidine has been found in urine and fecal samples of mammals and chicken and hence, guanidine-degrading bacteria contribute to N cycling in agricultural soils and in wastewater treatment plants (12). Moreover, enzymes that can produce relevant quantities of guanidine exist in plants and other photosynthetic eukaryotes (72). By assimilating guanidine, cyanobacteria could also largely contribute to the reintroduction of free guanidine into the N cycle. Albeit the environmental concentrations of guanidine and its meaning for the biogeochemical N cycle have not been systematically quantified yet, significant amounts and a widespread occurrence of guanidine can be expected due to the widespread occurrence of guanidine-degrading enzymes and regulatory systems for its perception. This is supported by the fact that guanidine can be obtained from natural resources, e.g. by oxidative degradation of guano, the accumulated excrement of seabirds or bats as already described in 1861 (73). In habitats exposed to seabird guano, guanidine assimilation appears advantageous. Indeed, *Synechocystis* and *Synechococcus* sp. PCC 7002 are euryhaline cyanobacteria (74) that are predestined to thrive in environments with fluctuating salinities, found in coastal areas close to river estuaries. In contrast, *S. elongatus* lacking GdmH is a typical freshwater strain and does not tolerate salinities of marine environments (75) and consequently are not present there. However, there is no clear correlation between the presence or absence of GdmH and the habitat in which a cyanobacterial strain is found.

### Guanidine assimilation versus toxicity

There is a clear correlation between GdmH presence and growth on guanidine as demonstrated for two other strains in addition to *Synechocystis*. However, externally supplied guanidine was also found to be quite toxic in concentrations >5 mM. GdmH is essential to cope with these conditions as a corresponding deletion mutant of *Synechocystis* or strains that naturally lack a *gdmH* gene barely grow in the presence of even trace amounts of guanidine. Consequently, GdmH may have evolved as enzyme for guanidine assimilation, or for guanidine detoxification. However, the NtcA-dependent induction of GdmH and the guanidine ABC transporter under N starvation suggest that the system rather evolved as an alternative assimilation mechanism. It should be noted that the responses to guanidine differed distinctly between the *Synechocystis* Δ*gdmH* strain and *S. elongatus* PCC 7942, which naturally lacks a guanidine-degrading enzyme. In *Synechocystis* Δ*gdmH*, growth halts almost immediately after guanidine is added, even at low concentrations. In contrast, *S. elongatus* continues to grow for a period that inversely correlates with guanidine concentration. This suggests an alternative, though less efficient, guanidine tolerance mechanism in *S. elongatus*, possibly involving a guanidine-degrading or efflux system with limited capacity.

Guanidine toxicity has previously been noticed in a *S. elongatus* strain engineered to produce ethylene via an ethylene-forming enzyme (EFE). Ethylene production was found unstable due to the intercellular accumulation of the toxic by-product guanidine (14). Accordingly, co-expression of an *efe* gene with a *gdmH* homolog from *Synechocystis* resulted in stable ethylene production in *S. elongatus* line, emphasizing the role of GdmH in coping with and avoiding high intracellular guanidine concentrations (14). *Synechocystis* has also been engineered towards ethylene production using EFE variants (76, 77). Interestingly, during ethylene production significant upregulation has been reported for the genes *sll1077-1079* (*gdmH (speB2), hypA2, hypB*) in contrast to other NtcA-regulated genes such as *glnA* (78). This observation is consistent with a guanidine-induced up-regulation mediated by the guanidine riboswitch in addition to NtcA-induced transcription.

The increased guanidine tolerance in *Synechocystis* strains with a deregulated PrqA multidrug efflux pump, combined with a likely guanidine-specific, NtcA-regulated ABC transporter, suggests that *Synechocystis* actively regulates its internal guanidine concentration. This regulation allows the otherwise toxic guanidine to be utilized as a N source across a wider range of external concentrations. At low guanidine levels, the ABC transporter (encoded by *sll1080*-*sll1082*) actively imports guanidine, maintaining internal concentrations suitable for assimilation by GdmH. In contrast, at high external guanidine concentrations, the PrqA efflux pump helps preventing toxic accumulation by exporting excess guanidine, allowing for continued assimilation. The appearance of suppressor mutants after long-term cultivation in presence of toxic guanidine concentrations positively affecting guanidine export by PrqA also explain previous results showing that *Synechocystis* can grow slowly on 5 mM guanidine after 2-3 weeks (10). The complete assimilation of guanidine by *Synechocystis* in the presence of nitrate indicates that *gdmH* is primarily regulated by the guanidine riboswitch, aligning an additional role for GdmH in guanidine detoxification. Even at low guanidine concentrations, which require active import via the ABC transporter system, assimilation occurs rapidly under nitrate-excess conditions. The reported basal activity of P*_sll1080_* or potential read-through from P*_gdmH_* under these conditions may provide sufficient ABC-transporter capacity for this process.

The assimilation of otherwise toxic compounds is a widespread phenomenon giving a clear advantage to the respective strains in exposed environments. For instance, *Pseudomonas putida* DOT-T1E is capable of assimilating the toxic compound toluene as a carbon source using the toluene dioxygenase (TOD) pathway. Similar to *Synechocystis* behaviour towards guanidine, *Pseudomonas putida* DOT-T1E activates assimilatory genes at low concentrations, while simultaneously maintaining efflux systems for toxic concentrations of toluene (79). Moreover, it should be noted that ammonium, the preferred N source to be assimilated, is also toxic for oxygenic phototrophs including cyanobacteria at high concentrations as it triggers photodamage of photosystem II (80). The repair requires the protease FtsH2, which is responsible for maintaining the turnover of the D1 protein (81, 82). Accordingly, a *ftsH2* knockout strain of *Synechocystis* was not able to survive in medium containing 5 mM NH_4_Cl, even though ammonium is assimilated. Similarly, we observed that guanidine induces also a strong reduction in PSII activity at high concentrations, presumably due to ammonium accumulation through the GdmH activity, or directly through attacking photosystem reaction center. Altogether, a certain toxicity and a relevance as substrate for growth do not exclude each other.

### Guanidine as a cost-effective inducer for large scale applications

Although various riboswitch classes have been identified in cyanobacteria, only a few are biochemically validated (41). Among them are glutamine riboswitches that are also crucial to control N assimilation (47). Riboswitches are of great interest for metabolic engineering as they link gene expression to internal metabolite levels. However, only a few riboswitch designs are in common use for metabolic engineering applications in cyanobacteria (40). Moreover, these riboswitches, as well as most of the inducible promoters commonly employed in cyanobacteria rely on comparatively costly chemicals such as L-rhamnose or theophylline, which limits large-scale commercial application (for price comparison see **Supplementary Table S1**). The here provided series of inducible expression systems building on the guanidine riboswitch provides an elegant solution. In GdmH^+^ strains, they allow rather inexpensive induction with a broad range of induction strength, titratability, and low leakiness. In contrast to the even cheaper metal inducers Ni^2+^ or Cu^2+^, guanidine likely exhibits lower toxicity at concentrations needed for full induction and better long-term activity. This is because guanidine is actively imported via the ABC-transporter system rather than being exported as the heavy metals (83). Unlike many heterologous inducible promoters, riboswitches are functional without the coincidental expression of a specific proteinaceous regulator. In some cases, these regulators need to be expressed from strong promoters to ensure tight control. Therefore, the additional metabolic load and potential pleiotropic effects can be avoided using riboswitches. Moreover, guanidine is biologically degradable avoiding elaborate removal of heavy metal ions from wastewater. Furthermore, the assimilation of guanidine as N source by *Synechocystis* allows its dual use as both N source and inducer of gene expression further cutting the induction-specific costs in a biotechnological application. Switching the N source to guanidine can simultaneously induce target genes, as demonstrated in the BVMO-mediated conversion of cyclohexanone to ε-caprolactone in a steady state system. Using guanidine in such scenario might also be advantageous for keeping producer strains axenic due to its toxicity as long as efficient guanidine-degrading mechanisms are absent. For other applications, such as CRISPR/Cas mediated genome editing or expression of toxic genes, the use of a riboswitch to control gene expression at mRNA level allows its combination with controllable promoters as shown here with the rhamnose-inducible system. This enables tight regulation and prevents potentially toxic or unfavorable basal expression.

## Conclusion

Unlike typical detoxification mechanisms that export or modify toxic substances, the GdmH allows a variety of cyanobacterial strains to exploit the otherwise toxic guanidine as a nutrient. So far, the occurrence and concentration of guanidine in natural environments is underexplored, but (at least) temporary and localized occurrence of relevant concentrations are to be expected and confirmed. In such environments, the evolution of guanidine hydrolase activity could have offered a strong selective advantage of detoxification by simultaneously releasing ammonium as a utilizable N-source further expanding its benefit. Together with the urea-metabolizing system, GdmH enabled complete utilization of guanidine as source of three N and a C and thereby the capacity for guanidine-supported growth instead of toxicity. The identification of a guanidine ABC transporter system expands the guanidine metabolism in order to fully exploit this potential by selectively importing the previously toxic substance. Despite these comprehensive benefits of guanidine assimilation, the intracellular concentration appears to be under control of importer as well as exporter systems balancing its benefit as a nutrient and harm as a toxin.

## Material and Methods

### Strains and growth conditions

*Escherichia coli* (*E. coli*) strains DH5α or TOP10 were cultivated on agar-solidified lysogeny broth (LB) medium or in liquid under constant shaking at 200 rpm. Depending on the carried plasmid, the strains were incubated at 37°C (pBR322-based plasmids and pSEVA351) or 30°C (pSOMA17). For selection of recombinant strains, the medium was supplemented with 100 µg/ml ampicillin (Amp), 50 µg/mL kanamycin (Kan), 100 µg/mL spectinomycin (Spec), 15 µg/mL chloramphenicol (Cm), or 10 µg/mL gentamycin (Gent).

*Synechocystis* sp. PCC 6803 and *Synechococcus elongatus* PCC 7942 were obtained from the Pasteur Culture Collection of Cyanobacteria (PCC) and served as wildtype (WT). *Synechococcus* sp. PCC 7002 was received from the lab of M. Hagemann (University of Rostock, Germany). By default, cyanobacterial strains were cultivated on plates using BG11 medium solidified with 1.5 % (w/v) Bacto agar (Becton Dickinson), buffered to pH 8.0 with 10 mM HEPES and supplemented with 0.3 g/L Na_2_S_2_O_3_. For liquid cultures, BG11 medium with the following modifications compared to the original protocol (84) was used: ferric ammonium citrate was replaced with 0.97 mg/L FeCl_3_, Na_2_EDTAx2H_2_O concentration was elevated to 5.95 mg/L, citric acid was omitted, medium was buffered to pH 8.0 with 10 mM HEPES. For the cultivation of *Synechococcus* sp. PCC 7002 the BG11 medium was additionally supplemented with 4 µg/L vitamin B12. For the cultivation on different N sources, we used a BG11 medium free of any N source (BG11^0^) which was then supplemented with NaNO_3_ or guanidine hydrochloride at the concentrations given for each experiment separately. If not stated otherwise, photoautotrophic culture conditions were set to constant illumination at 25 or 50 µmol_photons_ m^−2^ s^−1^, ambient CO_2_, 70 % (v/v) humidity, and a temperature of 30°C. Liquid cultures were shaken at 150 rpm in a Multitron Pro (INFORS HT), while agar plates were incubated in a SE41-CU5CLT growth chamber (CLF Plantclimatics). Mutants of *Synechocystis* sp. PCC 6803 were cultivated in presence of 50 µg/mL Kan (Δ*gdmH*; Δ*sll1080*; Δ*sll1080/81*) and 50 µg/mL Spec (Δ*sll1081*; Δ*sll1080/81*) as well as 15 µg/mL Cm when transformed with pSEVA351 derivatives or 5 µg/mL Gent when transformed with pSOMA17 plasmids.

In experiments in which strains with different antibiotic resistances were compared or growth performance was analyzed, antibiotics were omitted. Growth performances were either recorded in 250 mL Erlenmeyer shaking flasks with 25 mL culture volume by sampling and evaluating optical density at 750 nm (OD_750_) using a VIS-Spectrometer Libra S11 (biochrom). Alternatively, OD was automatically monitored in 10 min intervals from air-bubbled culture columns at 720 nm in Multicultivator® MC1000 reactors (Photon Systems Instruments). For steady state cultures of *Synechocystis*, the Multicultivator® MC1000 was additionally equipped with turbidostat modules allowing OD-based feeding and external peristaltic pumps to remove surplus culture volume.

In case of the drop dilution assays, cells from a rapidly growing liquid culture (OD_750_ < 1) were harvested by centrifugation (5 min, 3,900 x g), washed in 1 mM HEPES (pH 7.4), and re-suspended in 1 mM HEPES (pH 7.4) to a final OD_750_ of 0.5. From these suspensions, a 10-fold serial dilution with 4 steps (10^0^-10^−4^) in 1 mM HEPES (pH 7.4) was prepared. From each dilution of each analyzed strain 5 µl volume was dropped on a BG11 agar plate, which was incubated as stated above.

### Generation of genetic constructs to obtain recombinant Synechocystis strains

Genetic constructs were obtained by standard molecular cloning procedures using polymerase chain reaction (PCR) with Phusion polymerase, FastDigest restriction endonucleases and T4 DNA ligase (obtained from Thermo Scientific, New England Biolabs and/or Promega) by following the manufacturer’s instructions. PCR products were either cloned into pMiniT 2.0 using a PCR cloning kit (New England Biolabs) or used directly in assembly reactions. Plasmids were maintained in *E. coli* DH5α or *E. coli* TOP10. Final constructs were verified by Sanger sequencing (Azenta Life Sciences). Information about used oligonucleotides and plasmids is given in **Supplementary Tables S2** and **S3**. In addition, a previously introduced Golden-Gate cloning (GGC) system was used to assemble several plasmids in this study. The respective GGC library (85) was expanded by some standard parts and new entry plasmids. Their generation is described in the Appendix.

The *gdmH* gene (Gene ID: *sll1077*) was deleted via homologous recombination. For this, the chromosomal regions up- and downstream of the *gdmH* gene were amplified by PCR from genomic DNA (gDNA) of *Synechocystis* using primer pairs P1/P2 and P3/P4. Subsequently, the PCR products were fused in a scarless way during a BpiI-mediated Golden Gate assembly (GGA) into pGGC 78 yielding pAI 132. The fused up- and downstream region was then used to insert a kanamycin resistance cassette between the upstream located gene *slr1142* and the promoter region and RF00442 fused to *hypA2.* For this insertion, two PCR products were produced using primer pairs P1/P6 and P4/P5 with pAI 132 as template. The resulting products were fused with a Kan^R^::T*tonB* cassette from pGGC 335 into the level P entry vector pGGC 78 yielding pAI 140.

In a similar way, plasmids were generated to establish a disruption within genes *sll1080* and *sll1081*. The homologous regions were amplified using primer pairs P9/P10 and P11/P12 and subsequently inserted into pGGC 0 yielding pGGC 132 and pGGC 133. The level 0 plasmids of the homologous regions for *sll1081* interruption - pGGC 134 and pGGC 135 – were made with PCR products using primer pairs P13/P14 and P15/P16. In a second step, the upstream regions were transferred from their level 0 plasmid into the level 1 position 6 backbone pGGC 6 yielding pGGC 138 and pGGC 203, while the downstream regions were introduced into the level 1 position 4 backbone pGGC 4 resulting in pGGC 136 and pGGC 137. From there, the up- and downstream regions for the *sll1080* interruption were combined on the pBluescript level 2 plasmid pGGC 48 with the Kanamycin resistance cassette from pGGC 90 and the endlinker sequences from pGGC 139 and pGGC 47 generating the recombination template pGGC 204. The recombination template for *sll1081* interruption pGGC 205 was produced similarly by fusing the up- and downstream regions from pGGC 203 and pGGC 137 with the Spec^R^ cassette from pGGC 22 and the endlinkers from pGGC 139 and pGGC 47 on the pBluescript backbone pGGC 48.

Fluorescent guanidine reporter strains maintained pSEVA351 derivatives with a *sfgfp* gene that was transcriptionally fused to a constitutive promoter and the guanidine I riboswitch. For this the 5’UTR of *sll1077* including the RNA motif RF00442 was amplified from *Synechocystis* gDNA using primer pairs P17/P19 (WT variant) or P18/P19 (mutated variant, G at position 16 was replaced by C). Via the primers, the promoter P_J23101_ sequence was introduced upstream of RF00442. The *sfgfp* reporter gene was amplified from pXG10_SF (86) using the primer pair P20/P21. Due to overlapping sequences included in P19 and P20, both the *sfgfp* element and the 5’UTRs were fused and re-amplified in a second PCR using the primer pairs P17/P21 and P18/P21, respectively. Both fusion products were inserted into the multi cloning site of pSEVA351 using KpnI and EcoRI yielding pSEVA351-P_J23101_::RF00442^WT^::*sfgfp or* pSEVA351-P_J23101_::RF00442^G16C^::*sfgfp*, respectively. Analogously, the promoter of gene *sll1080* (Psll1080) was fused to *sfgfp*. For this, the region upstream of *sll1080* was amplified from *Synechocystis* gDNA using primer combinations P60/P61 or P62/P61 to obtain the native or a mutated sequence variant, respectively. The *sfgfp* gene was amplified from pXG10_SF with primer pair P63/P64. The Psll1080::*sfgfp* or Psll1080_mut::*sfgfp* constructs were maintained on the pSEVA351 derivatives pGGC 326 and pGGC 327, respectively. The generation of the plasmid series featuring the guanidine riboswitch is described in detail in **Supplementary Table S4**. In brief, the plasmids were constructed in a modular GGA approach in which the respective promoter was fused to the riboswitch, either *sfgfp* or a multi-cloning-site (MCS), and the terminator T_psbC_ on a pSEVA351 or pSOMA17 vector.

To enable an insertion of the *bvmo* gene from *Acidovorax* sp. CHX100 into the MCS of pAI234, the gene was flanked with PstI and EcoRI restriction sites via a PCR from pGGC 199 using primer pair P54/P55 and subsequently introduced into pGGC 0 using a BsaI-based GGA yielding pAI237. From there, the *bvmo* gene was transferred into the PstI/EcoRI sites of pAI234 using restriction and ligation giving access to the final plasmid pAI243.

Recombinant *Synechocystis* cells harbouring chromosomal changes were obtained via homologous recombination after incubating naturally competent cells with the corresponding plasmids and subsequent selection by antibiotics. The introduction of replicative plasmids by electroporation has been described previously (63, 87).

### Genomic DNA extraction and whole-genome sequencing of suppressor mutants

Genomic DNA was extracted from *Synechocystis* following standard protocols. Briefly, cells were treated with lysozyme (5 mg/mL) and RNase A (0.1 μg/mL) at 37 °C for 1 hour, followed by proteinase K (2 μL/mL) and SDS (2 % final) incubation at 60 °C overnight. DNA was extracted and purified by mixing several times with phenol/chloroform/isoamyl alcohol and chloroform followed by centrifugation (14000 x g for 10 min at 4 °C) and collecting the supernatants each time. DNA was precipitated with isopropanol, and the DNA pellets were washed with 70 % (v/v) ethanol, air-dried, and resuspended in nuclease-free water. Isolated genomic DNA of the guanidine resistant strains was sent to GENEWIZ (Leipzig, Germany) for library preparation and sequencing using an Illumina 2×150 bp configuration. Bioinformatic analysis was performed by GENEWIZ to assemble the genome of the guanidine resistant strains versus the published genome of *Synechocystis* PCC 6803. The mutations in the *prqR* (*slr0895*) were confirmed using the Integrative Genomics Viewer software.

### Riboswitch construct preparation and in-line probing

DNA oligonucleotides (**Supplementary Table S2**) were ordered from Microsynth AG and used to generate *in vitro* transcription templates by overlap extension PCR. For this, 400 nM of the two partially overlapping oligonucleotides, 1x Phusion HF buffer (Thermo Fisher), 150 µM dNTPs, 2 U Phusion polymerase (Thermo Fisher) and dH_2_O were mixed in 200 µl. The template was purified by phenol-chloroform extraction and ethanol precipitation (88). For subsequent *in vitro* transcription, 50 U T7 polymerase, 4.2 µM NTPs, 20 mM MgCl_2_, 40 mM DTT, 2 mM spermidine, ~ 0.5 µM purified DNA template and 3 % DMSO and DEPC-treated water were mixed, incubated at 37 °C overnight and purified by denaturing polyacrylamide gel electrophoresis (PAGE). Following *in vitro* transcription, the RNA was 5 [labeled using 10 U T4 PNK (NEB) and 50 pmol of radioactively labeled [γ-^32^P]-ATP in 1x T4 PNK reaction buffer (NEB) and DEPC-treated water. The total reaction volume was 20 µl and samples were incubated at 37 °C for 1 h before purification by denaturing 10 % (8 M urea) PAGE. In-line probing (ILP) was performed as previously described in (61) with minor alterations: RNA was heated in DEPC-treated H_2_O and in the presence of the ligand for 1 min at 90 °C, 2 min at 65 °C and 15 min at 37 °C. ILP buffer was added to start the ILP reaction. After 48 h at 23 °C, ILP reactions were quenched by adding 2x stop solution (90 % formamide and 50 mM EDTA), separated by denaturing (8 M urea) PAGE and visualized on a phosphorimager (cytiva). Guanidine hydrochloride was purchased from Sigma Aldrich. For K_D_ analysis, band intensities were determined with ImageQuant TL v8.1 and further calculations were performed in Microsoft Excel and GraphPad Prism.

### GFP fluorescence detection and microscopy imaging

*In vivo* GFP fluorescence was determined at excitation/emission wavelengths 485/520 nm from *Synechocystis* cells grown under standard conditions and in presence of different guanidine concentrations, in each case added as guanidine-HCl. Prior to fluorescence measurement, the cell samples were diluted to an OD_750_ of 0.2 and a volume of 200 µL was transferred to opaque black flat microtiter 96-well plates (Nunc). GFP fluorescence measurements were performed in an Infinite 200 PRO microplate reader as described previously (63). Final fluorescence intensity calculation involved background subtraction, normalization to corresponding OD_750_ values monitored in parallel and averaging of three biological replicates (= independently obtained clones), each measured in technical triplicates.

For fluorescence microscopy, cells were immobilized on an agarose pad, transferred onto a 24 x 60 mm high precision microscope cover glass (Zeiss, 1705µm, No. 1.5H) and visualized using a Zeiss AxioObserver.Z1/7 microscope with the Plan-Apochromat 100x/1.40 Oil Ph 3 M27 objective lens (Zeiss). GFP was detected with a filter set to 450 – 490 nm (ex.) and 500 – 550 nm (em.). Autofluorescence was measured at excitation/emission wavelengths 599/625 nm. Images were processed using ZEN 3.8 (ZEN lite).

### Metabolite analytics

To monitor guanidine uptake of the guanidine-supplemented cultures via HPLC, 1 mL of cell suspension was centrifuged at 17,000 g for 10 min and the supernatant was transferred into 2 ml threaded glas vials ND8 (LABSOLUTE). The analysis was performed using the UltiMate^TM^ 3000 system (Thermo Fisher) equipped with a Primesep 100 column (100Å 5µm, 4.6 x 250mm, Sielc Company). A total of 10 µl of sample was injected, and analytes were separated at 30°C oven temperature with an isocratic gradient consisting of 75 % water mixed with 0.25 % TFA (Trifluoroacetic acid) and 25 % ACN (acetonitrile) for 14 min. For detection a CAD (charged aerosol detector) was used with 60°C evaporator temperature, 20 Hz data collection rate and a filter value of 10.

To analyze the concentration of cyclohexanone and ε-caprolactone, both substances were extracted from the cell culture sample as described (63) and subsequently analyzed by gas chromatography analogous to previous work (89).

### Pulse-Amplitude-Modulation (PAM) Fluorometry

The photosystem II (PSII) activity was measured using a WATER-PAM-II chlorophyll fluorometer (Walz GmbH), as described previously (90). Samples were diluted in water to a final volume of 2 mL (ensuring no visible coloration) and normalized to a baseline fluorescence (F) between 400 and 500. Before measurement, samples were dark-adapted for 2 minutes. The first measurement was omitted, and the subsequent three were recorded as technical triplicates at 30-second intervals. Measurements were conducted using the saturation pulse method with a measuring light wavelength of 650 nm. The PSII activity was determined using the F_v_/F_m_ ratio, where F_v_ (Variable Fluorescence) = F_m_ (Maximal Fluorescence) − F_0_ (Minimal Fluorescence).

### Computational analyses

The software Geneious (Biomatters) was used to perform *in silico* molecular biology work. The phylogenetic tree based on 16S rDNA sequences of 35 cyanobacterial strains was generated by using the Maximum Likelihood method and Tamura-Nei model (91). Evolutionary analyses were conducted in MEGA11 (92). Presence or absence of homologous sequences to the experimentally identified GdmH of *Synechocystis* was analyzed using the BlastP algorithm and a part of a previously identified consensus motif for guanidine hydrolases (used amino acid sequence: SFDIDCIDAGFVPGTGWPEPGGLLPREAL, extracted from (10)). Synteny analysis was performed using the Integrated Microbial Genomes & Microbiomes (IMG/MER) database. A steady-state bioprocess was simulated using differential equations that were numerically solved using the software Berkeley Madonna (version 8.3.18). The employed code is given in the supplementary information.

## Acknowledgements

The UFZ is supported by the European Regional Development Funds (EFRE, Europe funds Saxony) and the Helmholtz Association. This research was funded by the Helmholtz Association and the Deutsche Forschungsgemeinschaft (individual grants KL 3114/7-1 and KL 3114/10-1 to S.K.; and SE 3449/1-1, SE 3449/3-1 and SFB1381-project number: 4032227 to K.A.S). We also acknowledge Wolfgang R. Hess and Karl Forchhammer for critically reading the manuscript, Rocio López-Igual for sharing materials, and Adrian Tüllinghoff for reviewing the code used for process simulations.

## Author Contributions

S.K. planned and designed the study; M.A.I., A.M.E., R.St., H.H., R.Sc. and L.M.B. performed research; C.E.W. designed and J.A.D. performed *in vitro* guanidine binding experiments; K.A.S., C.E.W. and S.K. supervised research, analyzed and interpreted data; and S.K. wrote the paper with contributions from all co-authors.

## Competing Interest Statement

The authors declare to have no significant competing financial, professional, or personal interests that might have influenced the performance or presentation of the work described in this manuscript.

## Notes

### Competing Interest Statement

The authors have declared no competing interest.

## References

1. L. Reitzer, Nitrogen assimilation and global regulation in *Escherichia coli*. Annu. Rev. Microbiol. 57, 155–176 (2003).

2. K. Forchhammer, K. A. Selim, Carbon/nitrogen homeostasis control in cyanobacteria. FEMS Microbiol. Rev. 44, 33–53 (2020).

3. R. H. H. van den Heuvel, B. Curti, M. A. Vanoni, A. Mattevi, Glutamate synthase: a fascinating pathway from L-glutamine to L-glutamate. Cell. Mol. Life Sci. CMLS 61, 669– 681 (2004).

4. E. Flores, J. E. Frías, L. M. Rubio, A. Herrero, Photosynthetic nitrate assimilation in cyanobacteria. Photosynth. Res. 83, 117–133 (2005).

5. J. T. Lin, V. Stewart, Nitrate assimilation by bacteria. Adv. Microb. Physiol. 39, 1–30, 379 (1998).

6. H. L. Mobley, R. P. Hausinger, Microbial ureases: significance, regulation, and molecular characterization. Microbiol. Rev. 53, 85–108 (1989).

7. T. Kanamori, N. Kanou, H. Atomi, T. Imanaka, Enzymatic characterization of a prokaryotic urea carboxylase. J. Bacteriol. 186, 2532–2539 (2004).

8. A. A. Kermani, C. B. Macdonald, R. Gundepudi, R. B. Stockbridge, Guanidinium export is the primal function of SMR family transporters. Proc. Natl. Acad. Sci. U. S. A. 115, 3060– 3065 (2018).

9. J. W. Nelson, R. M. Atilho, M. E. Sherlock, R. B. Stockbridge, R. R. Breaker, Metabolism of free guanidine in bacteria is regulated by a widespread riboswitch class. Mol. Cell 65, 220– 230 (2017).

10. D. Funck, et al., Discovery of a Ni^2+^-dependent guanidine hydrolase in bacteria. Nature 603, 515–521 (2022).

11. M. Sinn, F. Hauth, F. Lenkeit, Z. Weinberg, J. S. Hartig, Widespread bacterial utilization of guanidine as nitrogen source. Mol. Microbiol. 116, 200–210 (2021).

12. M. Palatinszky, et al., Growth of complete ammonia oxidizers on guanidine. Nature 633, 646–653 (2024).

13. N. O. Schneider, et al., Solving the conundrum: Widespread proteins annotated for urea metabolism in bacteria are carboxyguanidine deiminases mediating nitrogen assimilation from guanidine. Biochemistry 59, 3258–3270 (2020).

14. B. Wang, et al., A guanidine-degrading enzyme controls genomic stability of ethylene-producing cyanobacteria. Nat. Commun. 12, 5150 (2021).

15. N. Nelson, A. Ben-Shem, The complex architecture of oxygenic photosynthesis. Nat. Rev. Mol. Cell Biol. 5, 971–982 (2004).

16. M. F. Hohmann-Marriott, R. E. Blankenship, Evolution of photosynthesis. Annu. Rev. Plant Biol. 62, 515–548 (2011).

17. T. W. Lyons, C. T. Reinhard, N. J. Planavsky, The rise of oxygen in Earth’s early ocean and atmosphere. Nature 506, 307–315 (2014).

18. R. M. Soo, J. Hemp, D. H. Parks, W. W. Fischer, P. Hugenholtz, On the origins of oxygenic photosynthesis and aerobic respiration in cyanobacteria. Science 355, 1436–1440 (2017).

19. D. G. Capone, J. P. Zehr, H. W. Paerl, B. Bergman, E. J. Carpenter, *Trichodesmium*, a globally significant marine cyanobacterium. Science 276, 1221–1229 (1997).

20. J. P. Montoya, et al., High rates of N_2_ fixation by unicellular diazotrophs in the oligotrophic Pacific Ocean. Nature 430, 1027–1032 (2004).

21. J. P. Zehr, et al., Unicellular cyanobacteria fix N_2_ in the subtropical North Pacific Ocean. Nature 412, 635–638 (2001).

22. T. H. J. Niedermeyer, Anti-infective natural products from cyanobacteria. Planta Med. 81, 1309–1325 (2015).

23. J. K. Nunnery, E. Mevers, W. H. Gerwick, Biologically active secondary metabolites from marine cyanobacteria. Curr. Opin. Biotechnol. 21, 787–793 (2010).

24. R. Reher, et al., Native metabolomics identifies the rivulariapeptolide family of protease inhibitors. Nat. Commun. 13, 4619 (2022).

25. D. C. Ducat, J. C. Way, P. A. Silver, Engineering cyanobacteria to generate high-value products. Trends Biotechnol. 29, 95–103 (2011).

26. N. M. Schmelling, M. Bross, What is holding back cyanobacterial research and applications? A survey of the cyanobacterial research community. Nat. Commun. 15, 6758 (2024).

27. J. Toepel, R. Karande, S. Klähn, B. Bühler, Cyanobacteria as whole-cell factories: current status and future prospectives. Curr. Opin. Biotechnol. 80, 102892 (2023).

28. K. Shabestary, et al., Design of microbial catalysts for two-stage processes. Nat. Rev. Bioeng. 1–17 (2024). 10.1038/s44222-024-00225-x.

29. M. J. Merrick, R. A. Edwards, Nitrogen control in bacteria. Microbiol. Rev. 59, 604–622 (1995).

30. I. Luque, E. Flores, A. Herrero, Molecular mechanism for the operation of nitrogen control in cyanobacteria. EMBO J. 13, 2862–2869 (1994).

31. A. Herrero, A. M. Muro-Pastor, E. Flores, Nitrogen control in cyanobacteria. J. Bacteriol. 183, 411–425 (2001).

32. J. Giner-Lamia, et al., Identification of the direct regulon of NtcA during early acclimation to nitrogen starvation in the cyanobacterium Synechocystis sp. PCC 6803. Nucleic Acids Res. 45, 11800–11820 (2017).

33. M. García-Domínguez, J. C. Reyes, F. J. Florencio, NtcA represses transcription of gifA and gifB, genes that encode inhibitors of glutamine synthetase type I from Synechocystis sp. PCC 6803. Mol. Microbiol. 35, 1192–1201 (2000).

34. P. Bolay, et al., The novel PII-interacting protein PirA controls flux into the cyanobacterial ornithine-ammonia cycle. mBio 12 (2021).

35. A. Kraus, et al., Protein NirP1 regulates nitrite reductase and nitrite excretion in cyanobacteria. Nat. Commun. 15, 1911 (2024).

36. M. I. Muro-Pastor, J. C. Reyes, F. J. Florencio, Cyanobacteria perceive nitrogen status by sensing intracellular 2-oxoglutarate levels. J. Biol. Chem. 276, 38320–38328 (2001).

37. J. Espinosa, K. Forchhammer, S. Burillo, A. Contreras, Interaction network in cyanobacterial nitrogen regulation: PipX, a protein that interacts in a 2-oxoglutarate dependent manner with P_II_ and NtcA. Mol. Microbiol. 61, 457–469 (2006).

38. K. Forchhammer, K. A. Selim, L. F. Huergo, New views on PII signaling: from nitrogen sensing to global metabolic control. Trends Microbiol. S0966-842X(21)00321–8 (2022). 10.1016/j.tim.2021.12.014.

39. K. A. Selim, M. Haffner, B. Watzer, K. Forchhammer, Tuning the in vitro sensing and signaling properties of cyanobacterial PII protein by mutation of key residues. Sci. Rep. 9, 18985 (2019).

40. M. Santos-Merino, A. K. Singh, D. C. Ducat, New applications of synthetic biology tools for cyanobacterial metabolic engineering. Front. Bioeng. Biotechnol. 7 (2019).

41. S. Klähn, F. Opel, W. R. Hess, Customized molecular tools to strengthen metabolic engineering of cyanobacteria. Green Carbon 2, 149–163 (2024).

42. M. Mandal, R. R. Breaker, Gene regulation by riboswitches. Nat. Rev. Mol. Cell Biol. 5, 451–463 (2004).

43. W. C. Winkler, R. R. Breaker, Regulation of bacterial gene expression by riboswitches. Annu. Rev. Microbiol. 59, 487–517 (2005).

44. J. E. Barrick, R. R. Breaker, The distributions, mechanisms, and structures of metabolite-binding riboswitches. Genome Biol. 8, R239 (2007).

45. M. Mandal, et al., A glycine-dependent riboswitch that uses cooperative binding to control gene expression. Science 306, 275–279 (2004).

46. A. Serganov, L. Huang, D. J. Patel, Structural insights into amino acid binding and gene control by a lysine riboswitch. Nature 455, 1263–1267 (2008).

47. S. Klähn, et al., A glutamine riboswitch is a key element for the regulation of glutamine synthetase in cyanobacteria. Nucleic Acids Res. 46, 10082–10094 (2018).

48. M. E. Sherlock, S. N. Malkowski, R. R. Breaker, Biochemical validation of a second guanidine riboswitch class in bacteria. Biochemistry 56, 352–358 (2017).

49. M. E. Sherlock, R. R. Breaker, Biochemical validation of a third guanidine riboswitch class in bacteria. Biochemistry 56, 359–363 (2017).

50. F. Lenkeit, I. Eckert, J. S. Hartig, Z. Weinberg, Discovery and characterization of a fourth class of guanidine riboswitches. Nucleic Acids Res. 48, 12889–12899 (2020).

51. H. Salvail, A. Balaji, D. Yu, A. Roth, R. R. Breaker, Biochemical validation of a fourth guanidine riboswitch class in bacteria. Biochemistry 59, 4654–4662 (2020).

52. K. Sidoruk, V. Melnik, M. Babykin, S. Shestakov, R. Cerff, “Cloning and Molecular Analysis of the Gene PRQR Controlling Resistance to Paraquat in Synechocystis sp. PCC 6803” in The Phototrophic Prokaryotes, G. A. Peschek, W. Löffelhardt, G. Schmetterer, Eds. (Springer US, 1999), pp. 715–718.

53. M. M. Babykin, et al., On the Involvement of the regulatory gene prqR in the development of resistance to methyl viologen in cyanobacterium Synechocystis sp. PCC 6803. Russ. J. Genet. 39, 18–24 (2003).

54. A. Klotz, et al., Awakening of a dormant cyanobacterium from nitrogen chlorosis reveals a genetically determined program. Curr. Biol. CB 26, 2862–2872 (2016).

55. S. E. Ongley, J. J. L. Pengelly, B. A. Neilan, A multidrug efflux response to methyl viologen and acriflavine toxicity in the cyanobacterium *Synechocystis* sp. PCC6803. J. Appl. Phycol. 28, 2793–2803 (2016).

56. R. I. Khan, et al., Transcriptional regulator PrqR plays a negative role in glucose metabolism and oxidative stress acclimation in *Synechocystis* sp. PCC 6803. Sci. Rep. 6, 32507 (2016).

57. A. Maqbool, et al., The substrate-binding protein in bacterial ABC transporters: dissecting roles in the evolution of substrate specificity. Biochem. Soc. Trans. 43, 1011–1017 (2015).

58. N. M. Koropatkin, H. B. Pakrasi, T. J. Smith, Atomic structure of a nitrate-binding protein crucial for photosynthetic productivity. Proc. Natl. Acad. Sci. U. S. A. 103, 9820–9825 (2006).

59. M. Kopf, et al., Comparative analysis of the primary transcriptome of *Synechocystis* sp. PCC 6803. DNA Res. 21, 527–539 (2014).

60. J. C. Reyes, M. I. Muro-Pastor, F. J. Florencio, Transcription of glutamine synthetase genes (*glnA* and *glnN*) from the cyanobacterium *Synechocystis* sp. strain PCC 6803 is differently regulated in response to nitrogen availability. J. Bacteriol. 179, 2678–2689 (1997).

61. E. E. Regulski, R. R. Breaker, In-line probing analysis of riboswitches. Methods Mol. Biol. Clifton NJ 419, 53–67 (2008).

62. D. Camsund, T. Heidorn, P. Lindblad, Design and analysis of LacI-repressed promoters and DNA-looping in a cyanobacterium. J. Biol. Eng. 8, 4 (2014).

63. F. Opel, et al., Generation of synthetic shuttle vectors enabling modular genetic engineering of cyanobacteria. ACS Synth. Biol. (2022). 10.1021/acssynbio.1c00605.

64. A. Behle, P. Saake, A. T. Germann, D. Dienst, I. M. Axmann, Comparative dose-response analysis of inducible promoters in cyanobacteria. ACS Synth. Biol. 9, 843–855 (2020).

65. M. J. L. J. Fürst, A. Gran-Scheuch, F. S. Aalbers, M. W. Fraaije, Baeyer–Villiger Monooxygenases: Tunable Oxidative Biocatalysts. ACS Catal. 9, 11207–11241 (2019).

66. A. Tüllinghoff, et al., Maximizing photosynthesis-driven Baeyer–Villiger oxidation efficiency in recombinant *Synechocystis* sp. PCC6803. Front. Catal. 1 (2022).

67. A. N. Muttathukattil, S. Srinivasan, A. Halder, G. Reddy, Role of guanidinium-carboxylate ion interaction in enzyme inhibition with implications for drug design. J. Phys. Chem. B 123, 9302–9311 (2019).

68. S.-H. Kim, D. Semenya, D. Castagnolo, Antimicrobial drugs bearing guanidine moieties: A review. Eur. J. Med. Chem. 216, 113293 (2021).

69. W. R. Mitchell, Microbial degradation of guanidinium ion. Chemosphere 16, 1071–1086 (1987).

70. C. Reiss, Y. Xiong, S. Strobel, Structural basis for ligand binding to the guanidine-I riboswitch. Struct. Lond. Engl. 1993 25, 195–202 (2017).

71. R. A. Battaglia, I. R. Price, A. Ke, Structural basis for guanidine sensing by the *ykkC* family of riboswitches. RNA 23, 578–585 (2017).

72. D. Funck, M. Sinn, G. Forlani, J. S. Hartig, Guanidine production by plant homoarginine-6-hydroxylases. eLife 12, RP91458 (2024).

73. A. Strecker, Untersuchungen über die chemischen Beziehungen zwischen Guanin, Xanthin, Theobromin, Caffein und Kreatinin. (1861). 10.1002/jlac.18611180203.

74. S. Klähn, M. Hagemann, Compatible solute biosynthesis in cyanobacteria. Environ. Microbiol. 13, 551–562 (2011).

75. M. Hagemann, Molecular biology of cyanobacterial salt acclimation. FEMS Microbiol. Rev. 35, 87–123 (2011).

76. F. Guerrero, V. Carbonell, M. Cossu, D. Correddu, P. R. Jones, Ethylene synthesis and regulated expression of recombinant protein in *Synechocystis* sp. PCC 6803. PloS One 7, e50470 (2012).

77. J. Ungerer, et al., Sustained photosynthetic conversion of CO_2_ to ethylene in recombinant cyanobacterium *Synechocysti* 6803. Energy Environ. Sci. 5, 8998–9006 (2012).

78. E. Kuchmina, et al., Ethylene production in *Synechocystis* sp. PCC 6803 promotes phototactic movement. Microbiol. Read. Engl. 163, 1937–1945 (2017).

79. T. Krell, et al., Responses of *Pseudomonas putida* to toxic aromatic carbon sources. J. Biotechnol. 160, 25–32 (2012).

80. M. Drath, et al., Ammonia triggers photodamage of photosystem II in the cyanobacterium *Synechocystis* sp. strain PCC 6803. Plant Physiol. 147, 206–215 (2008).

81. P. Silva, et al., FtsH is involved in the early stages of repair of photosystem II in *Synechocystis* sp PCC 6803. Plant Cell 15, 2152–2164 (2003).

82. J. Komenda, et al., The FtsH protease *slr0228* is important for quality control of photosystem II in the thylakoid membrane of *Synechocystis* sp. PCC 6803. J. Biol. Chem. 281, 1145–1151 (2006).

83. B. Blasi, L. Peca, I. Vass, P. B. Kós, Characterization of stress responses of heavy metal and metalloid inducible promoters in *Synechocystis* PCC6803. J. Microbiol. Biotechnol. 22, 166–169 (2012).

84. R. Rippka, J. Deruelles, J. B. Waterbury, M. Herdman, R. Y. Stanier, Generic assignments, strain histories and properties of pure cultures of cyanobacteria. J. Gen. Microbiol. 111, 1– 61 (1979).

85. S. Lupacchini, et al., Co-expression of auxiliary genes enhances the activity of a heterologous O_2_-tolerant hydrogenase in the cyanobacterium *Synechocystis* sp. PCC 6803. Biotechnol. Biofuels Bioprod. **accepted** (2025).

86. C. P. Corcoran, et al., Superfolder GFP reporters validate diverse new mRNA targets of the classic porin regulator, MicF RNA. Mol. Microbiol. 84, 428–445 (2012).

87. F. Brandenburg, et al., Trans-4-hydroxy-L-proline production by the cyanobacterium *Synechocystis* sp. PCC 6803. Metab. Eng. Commun. 12, e00155 (2021).

88. J. Sambrook, E. Fritsch, T. Maniatis, Molecular Cloning: A Laboratory Manual, 2nd edition (1989).

89. L. Schäfer, K. Bühler, R. Karande, B. Bühler, Rational engineering of a multi-step biocatalytic cascade for the conversion of cyclohexane to polycaprolactone monomers in *Pseudomonas taiwanensis*. Biotechnol. J. 15, e2000091 (2020).

90. K. A. Selim, et al., Diurnal metabolic control in cyanobacteria requires perception of second messenger signaling molecule c-di-AMP by the carbon control protein SbtB. Sci. Adv. 7, eabk0568 (2021).

91. K. Tamura, M. Nei, Estimation of the number of nucleotide substitutions in the control region of mitochondrial DNA in humans and chimpanzees. Mol. Biol. Evol. 10, 512–526 (1993).

92. K. Tamura, G. Stecher, S. Kumar, MEGA11: Molecular Evolutionary Genetics Analysis Version 11. Mol. Biol. Evol. 38, 3022–3027 (2021).

93. V. Krasikov, E. Aguirre von Wobeser, H. L. Dekker, J. Huisman, H. C. P. Matthijs, Time-series resolution of gradual nitrogen starvation and its impact on photosynthesis in the cyanobacterium *Synechocystis* PCC 6803. Physiol. Plant. 145, 426–439 (2012).

94. J. Mitschke, et al., An experimentally anchored map of transcriptional start sites in the model cyanobacterium *Synechocystis* sp. PCC6803. Proc. Natl. Acad. Sci. U. S. A. 108, 2124–2129 (2011).

