## Appendix for "When a chaotropic agent turns into a nutrient – Deciphering the assimilation of guanidine and its utilization to drive synthetic processes in cyanobacteria"

### Supplementary Figures and Tables

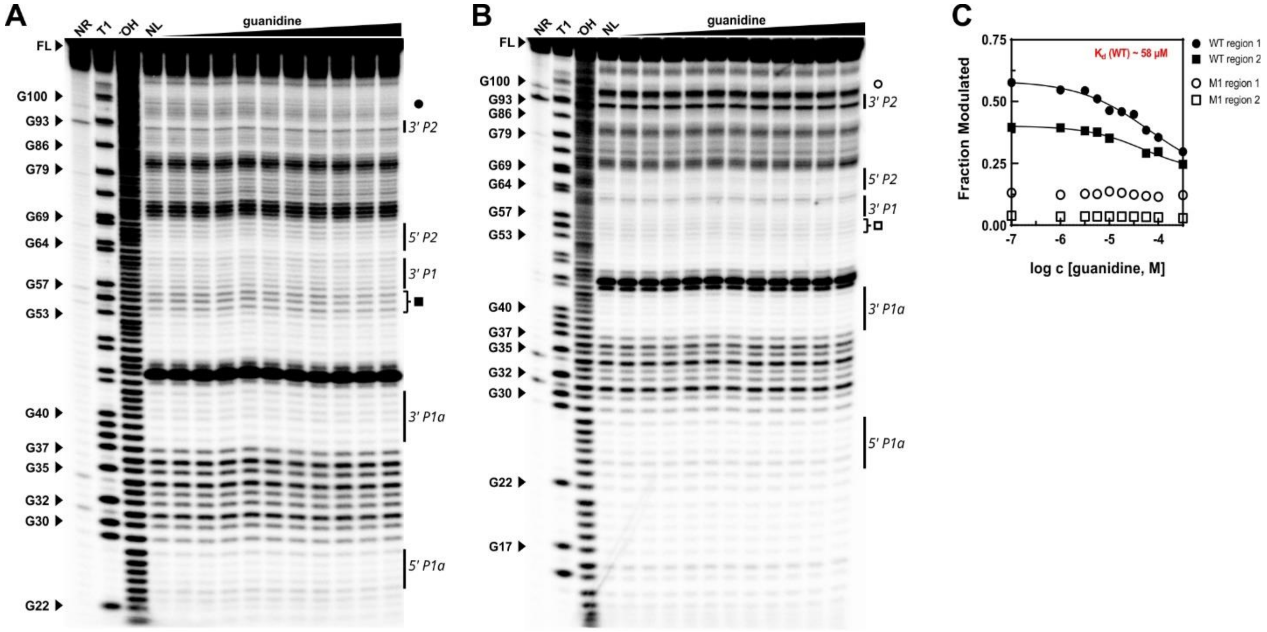


**Figure S1: Guanidine binding by the predicted aptamer in the untranslated region of the *gdmH* transcript. A:** In-line probing (ILP) analysis of RF00442-WT. Guanidine concentrations from 100 nM to 316 µM were tested. NR – no reaction control, T1 – RNase T1 digestion, $^{-}$OH – partial alkaline hydrolysis, NL – no ligand control. Regions with guanidine-dependent conformational changes are marked (Circle and Square). Note that the same gel as in Fig. 6A is shown here for comparison. **B:** ILP analysis of RF00442-M1 mutant. Annotations as in A) were used. RF00442-M1 showed no conformational changes upon incubation with guanidine. **C:** Graph of the $K_d$ analysis of RF00442-WT and – M1 using pixel analysis of band regions labeled with the corresponding symbols.

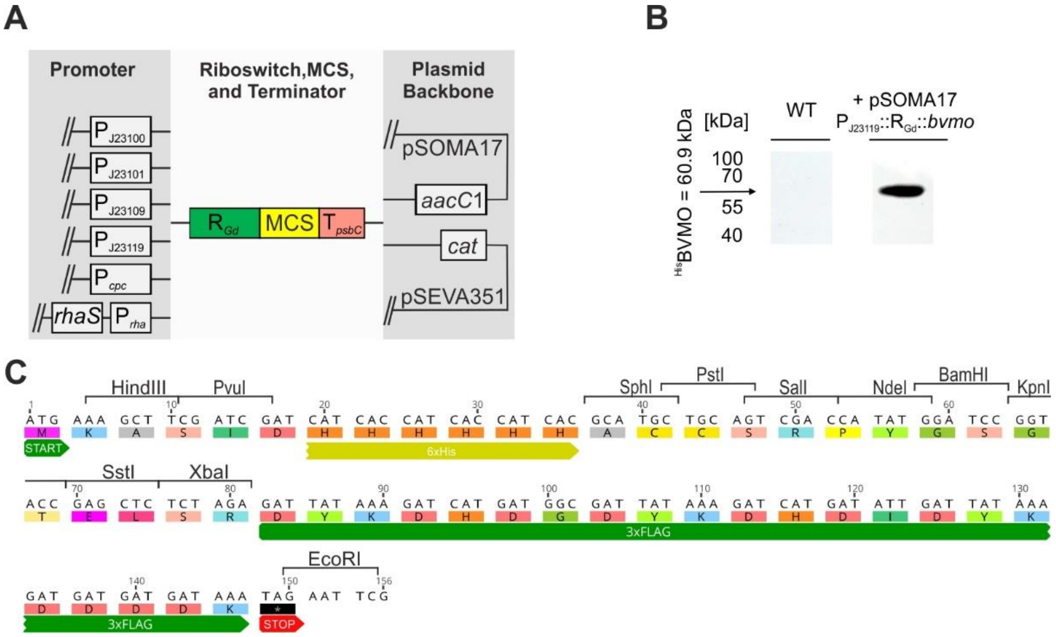


**Figure S2: Overview of the designed plasmid library and its possible applications.** **A:** Schematic overview of generated plasmid series based on two different plasmid backbones, pSOMA17 and pSEVA351 together with different promoters to drive transcription. In each case the guanidine riboswitch ($R_{Gd}$) is located downstream of the indicated promoters. A multiple cloning site (MCS) was added that enables integration of genes of interest via compatible restriction endonuclease sites. **B:** Western blot of proteins extracts from *Synechocystis* WT and a strain carying a plasmid with a *bvmo* gene inserted into MCS using a Anti-His-HRP conjugate antibody (Qiagen). **C:** Multiple cloning site in the generated vectors pAI225 – pAI235 and additional features that allow tagging of the introduced sequences.

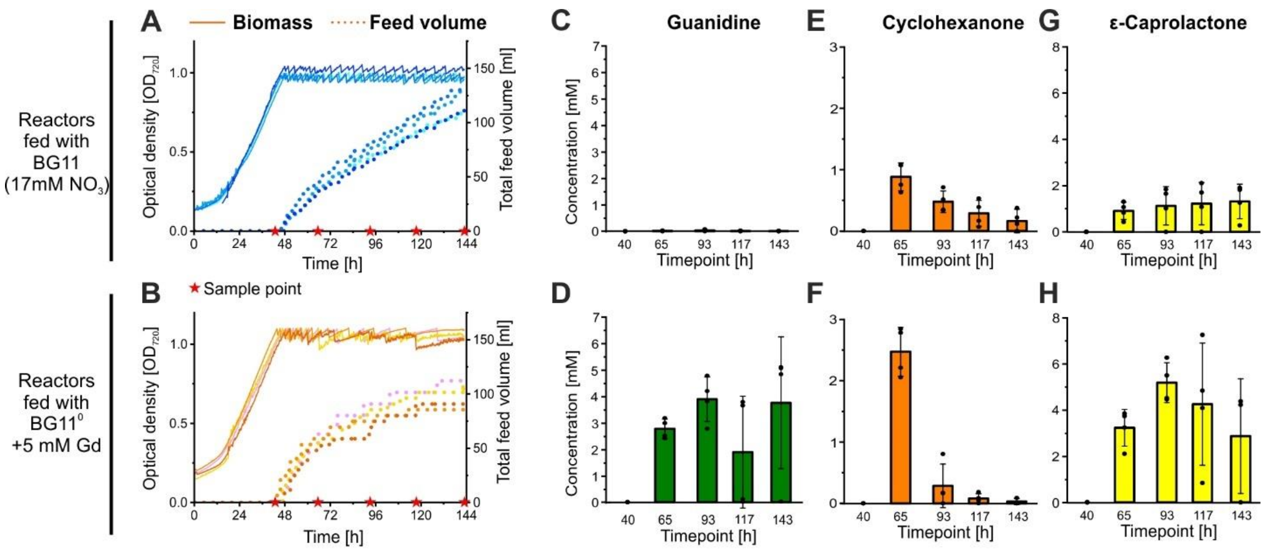


**Figure S3: Additional data of the continuous biotransformation of cyclohexanone to ε-caprolactone with guanidine as inducer of enzyme expression and N source.** Eigth cultures of 60 ml volume were managed in a steady state process using Multicultivators equipped with turbidostat modules. The cultures were inoculated to an $OD_{720}$ of ~ 0.1 in BG11 and grown to an $OD_{720}$ of 1 under bubbling with air enriched to 4% $CO_2$ and 200 µmol photons $m^{-2}$ $s^{-1}$**.** The biomass density was kept at an $OD_{720}$ of 1 by automatic, sequential dilution of the culture from the feed medium. The volume was kept constant by removing surplus culture via an external peristaltic pump. For four of the cultures (Panel A,C,E,G) the feed medium was BG11, while the other four cultures (Panel B,D,F,H) were fed with $BG11^0$ containing 5 mM guanidine as N-source instead of nitrate. Both feed-solutions were supplemented with 30 mM cyclohexanone as substrate for the biotransformation. Samples were taken from all cultures at the timepoints depicted as red stars and subsequently analyzed for their guanidine, cyclohexanone and ε-caprolactone concentration. **A,B**: Biomass density measured as $OD_{720}$ and total feed volume of the single cultures. **C,D:** Guanidine concentration in the samples shown as single points, average (green bar), and standard deviation (error bars). **E,F:** Cyclohexanone concentration in the samples shown as single points, average (orange bar), and standard deviation (error bars). G**,H:** ε-Caprolactone concentration in the samples shown as single points, average (orange bar), and standard deviation (error bars). Data from panel B and panel H is also shown in Fig. 8D and 8E.

**Table S1: Calculation of prizes for different chemical inducers used in cyanobacteria.**

| Inducer | Commonly used concentration | Amount needed for 1 $m^3$ | Prize per g | Supplier, date of access | Prize of Inducer for 1 $m^3$ | Source |
|---|---|---|---|---|---|---|
| Isopropyl β-D-thiogalactopyranoside (IPTG) | 1 mM | 238.3 g | 10.75€ | Carl Roth, 06.02.2025 | 2,562€ | (1) |
| Anhydrotetracycline | 10 mg/l | 10 g | 2.65€ | Carl Roth, 06.02.2025 | 27€ | (2) |
| 2,4-Diacetylphloroglucinol (DAPG) | 10 µM | 2.1 g | 229.95€ | Carl Roth, 06.02.2025 | 483€ | (3) |
| L-rhamnose | 10 mM | 1,821.7 g | 4.36€ | Carl Roth, 06.02.2025 | 7,943€ | (4) |
| L-arabinose | 10 mM | 1,501,3 g | 0.16€ | Carl Roth, 09.02.2025 | 243€ | (5) |
| $NiSO_4$ x 6 $H_2O$ | 10 µM | 2.6 g | 0.29€ | Carl Roth, 06.02.2025 | 0.75€ | (6) |
| CuCl | 3 µM | 0.3 g | 0.20€ | Carl Roth, 06.02.2025 | 0.06€ | (7) |
| Vanillic acid | 2 mM | 336.3 g | 1.28€ | Sigma Aldrich, 07.02.2025 | 430€ | (4) |
| Theophylline | 2 mM | 360.3 g | 0.30€ | Sigma Aldrich, 07.02.2025 | 106€ | (8) |
| Guanidine hydrochloride | 1.25 mM | 119.4 g | 0.05€ | Carl Roth, 06.02.2025 | 5.68€ | This study |

**Table S2: Sequence of primers used in this study.** The primer binding regions are given in bold, BpiI recognition sites are marked yellow, BsaI recognition sites are marked green, LguI recognition sites are marked cyan, recognition sites of Typ II restriction enzymes are marked red, cutting sites of the restriction enzymes are underlined, sequences of the promoter series $J_{23100}$ are given in yellow letters. Oligos used to generate *in vitro* transcription templates of the RF00442 riboswitch constructs are partially overlapping to be used in an overlap-extension PCR (OE) to synthesize the corresponding full-length constructs.

| **number** | **Primer name** | **Sequence 5´ -> 3´** | **Primer target, Purpose** |
|---|---|---|---|
| **P1** | 5'hom_gdmH for | AAAGAAGACAAGCCA**CTTCTAAAAACTGAACATCGTC** | slr1142 |
| **P2** | 5'hom_gdmH rev | AAAGAAGACAAG**CATAGGTAGTTACTAGCTAAAC** | 5'UTR of *gdmH* |
| **P3** | 3'hom_gdmH for | AAAGAAGACAA**ATGCATGAAACAGACATGACC** | *hypA2* |
| **P4** | 3'hom_gdmH rev | AAAGAAGACAAGATG**CTTCGGTGACACTGAGTAG** | *hypB* |
| **P5** | 3'hom_gdmH::AbR for | AAAGAAGACAAAAAG**TCGCCGATAAAAAGTAGCAC** | upstream region of *gdmH* |
| **P6** | 5'hom_gdmH::AbR rev | AAAGAAGACAACATC**TCGCCAAACTAAAGTTTAAAACC** | upstream region of *gdmH* |
| **P7** | KanR::TtonB for | AAAGGTCTCGGATG**CCGGAATTGCCAGCTGGG** | KanR |
| **P8** | KanR::TtonB rev | AAAGGTCTCGCTTT**AGTCAAAAGCCTCCGGTCG** | *tonB* terminator |
| **P9** | 5'hom_sll1080 for | AAAGGTCTCGGATG**GGCTAAGGCTTGCATGGATTC** | *sll1080* |
| **P10** | 5'hom_sll1080 rev | AAAGGTCTCGCTTT**CCAGGAAGCAGATTGTTTACTAATTAC** | *hypB* |
| **P11** | 3'hom_sll1080 for | AAAGGTCTCGGATG**CTCACCCCCGCCCGTTTAG** | *sll1080* |
| **P12** | 3'hom_sll1080 rev | AAAGGTCTCGCTTT**GCCGGGCAGTTGGATGG** | *sll1080* |
| **P13** | 5'hom_sll1081 for | AAAGGTCTCGGATG**GAAATCCATCCTTAAATAGCTTTATC** | *sll1081* |
| **P14** | 5'hom_sll1081 rev | AAAGGTCTCGCTTT**CTATTTAAGGATGGCACTCG** | *sll1080* |
| **P15** | 3'hom_sll1081 for | AAAGGTCTCGGATG**GAATCTGAGGATGAGGCGAAATG** | *sll1081* |
| **P16** | 3'hom_sll1081 rev | AAAGGTCTCGCTTT**TAATCCAGGATTCCCTGACTAGTTTTG** | *sll1081* |
| **P17** | EcoRI::PJ23101::$R_{Gd}$ for | TAGAATTCTTTACAGCTAGCTCAGTCCTAGGTATTATGCTAGC**ATATTGTTTCTAGGGTTCCGATTCC** | RF00442 |
| **P18** | EcoRI::PJ23101::$R_{Gd}$(M) for | TAGAATTCTTTACAGCTAGCTCAGTCCTAGGTATTATGCTAGC**ATATTGTTTCTAGCGTTCCGATTCC** | RF00442 |
| **P19** | RGd rev | CTTTGCTCAT**AGGTAGTTACTAGCTAAACAAC** | RF00442 |
| **P20** | sfGFP for | GTAACTACCT**ATGAGCAAAGGAGAAGAAC** | *sfgfp* |
| **P21** | KpnI::sfGFP rev | TAGGTACC**GGATTTGTCCTACTCAGGAG** | *sfgfp 3' UTR* |
| **P22** | LacZ_for | AAAACGCGT**GCGCAACGCAATTAATGTGAG** | lacZ fragment |
| **P23** | LacZ_rev | AAACCATGG**CTATGCGGCATCAGAGCAGA** | lacZ fragment |
| **P24** | BB_pUK21_LevP for | AAAACGCGTGAAGACTTTGGCTAACTGAGACC**ATTGCGTTGCGCTCACTG** | pUK21 |
| **P25** | BB_pUK21_LevP rev | AAACCATGGGAAGACAACATCTGTGAGAGACC**GGCGGGTGTGGTGGTTA** | pUK21 |
| **P26** | KanR for | AAAGGTCTCGGATG**CCGGAATTGCCAGCTGG** | Kan$^R$ |
| **P27** | KanR rev | AAAGGTCTCGCTTT**TCAGAAGAACTCGTCAAGAAG** | Kan$^R$ |
| **P28** | KanR+TtonB for | AAAGAAGACAAAGTCAAAAGCCTCCGACCGGAGGCTTTTGACT**AAAGGAGCAGAGACCGGCGGG** | Kan$^R$ |
| **P29** | KanR+TtonB rev | AAAGAAGACAAGACT**CAGAAGAACTCGTCAAGAAGGCGATAG** | Kan$^R$ |
| **P30** | CmcR for | AAAGGTCTCGGATG**TGATCGGCACGTAAGAGG** | Cmc$^R$ |
| **P31** | CmcR rev | AAAGGTCTCGCTTT**TTACGCCCCGCCCTGC** | Cmc$^R$ |
| **P32** | pBI-Backbone for | AAAGAAGACAAAAAG**CTGTCAGACCAAGTTTACTCATATATAC** | pGGC 46 |
| **P33** | pBI-Backbone rev | AAAGAAGACAACATC**TGTATTTAGAAAAATAAACAAATAGGGGTTC** | pGGC 46 |
| **P34** | P*cpc* for | AAAGGTCTCGGTTA**ACCTGTAGAGAAGAGTCCC** | Promoter of *cpc* |
| **P35** | P*cpc* rev | AAAGGTCTCGATAT**GGGATTTTCTTAAACACAATTCCC** | Promoter of *cpc* |
| **P36** | PJ23100 for | AAAGGTCTCGGTTATTGACGGCTAGCTCAGTCCTAGGTACAGTGCTAGC**ATATTGTTTCTAGGGTTCCG** | RF00442 |
| **P37** | PJ23101 for | AAAGGTCTCGGTTA**TTTACAGCTAGCTCAGTCC** | RF00442 |

| | | | |
|---|---|---|---|
| **P38** | PJ23109 for | AAAGGTCTCGGTTATTTACAGCTAGCTCAGTCCTAGGGACTGTGCTAGCATATTGTTTCTAGGGTTCCG | RF00442 |
| **P39** | PJ23119 for | AAAGGTCTCGGTTATTGACAGCTAGCTCAGTCCTAGGTATAATGCTAGCATATTGTTTCTAGGGTTCCG | RF00442 |
| **P40** | Prha for | AAAGGTCTCGGTTACCACAATTCAGCAAATTGTGAAC | Prha |
| **P41** | Prha rev | AAAGGTCTCGATATTTCATTACGACCAGTCTAAAAAGC | Prha |
| **P42** | RGd for | AAAGGTCTCGATATTGTTTCTAGGGTTCCG | RF00442 |
| **P43** | RGd rev 2 | AAAGGTCTCGTCATAGGTAGTTACTAGCTAAACAAC | RF00442 |
| **P44** | sfGFP for 2 | AAAGGTCTCGATGAGCAAAGGAGAAGAACTTTTC | *sfgfp* |
| **P45** | sfGFP rev | AAAGGTCTCGCTTTTTATTTGTAGAGCTCATCCATGC | *sfgfp* |
| **P46** | rhaS_Pos.1 for | AAAGGTCTCAGCCATTGACAGCTAGCTCAGTCC | *rhaS* |
| **P47** | rhaS_Pos.1 rev | AAAGGTCTCATAACATAAACGCAGAAAGGCCCAC | *rhaS* |
| **P48** | MCS for | AAAGGTCTCGATGAAAGCTTCGATCGATCATCACCATCACCATCACGCATGCTGCAGTCGACCATATGGATCCGGTACCGAGCTCTCTAGAGATTATAAA | - |
| **P49** | MCS rev | AAAGGTCTCGCTTTGCGAATTCTATTTATCATCATCATCTTTATAATCAATATCATGATCTTTATAATCGCCATCATGATCTTTATAATCTCTAGAGAGC | - |
| **P50** | pCB2.4 for | AAAGCTCTTCATACGTAGTTTAAACCGGCCG | *pCB2.4 backbone* |
| **P51** | pSOMA12/17 GGC rev | AAAGCTCTTCAAAATGGTCGCTTTCAGGGGTAC | *pSOMA12 and 17 backbone* |
| **P52** | lacZ cassette for | AAAGCTCTTCATTTCCGCCATGAGACCACGC | *BsaI recognition site flanking the lacZ casette* |
| **P53** | lacZ cassette rev | AAAGCTCTTCAGTAACCCGATGAGAGACCCTC | *BsaI recognition site flanking the lacZ casette* |
| **P54** | PstI-bvmo for | AAAGGTCTCGGATGCTGCAGTATGAAAAAAACCCAACATCTGG | *bvmo* |
| **P55** | EcoRI-bvmo rev | AAAGGTCTCGCTTTGAATTCTAAGCGCTCTGGAATACGAAAC | *bvmo* |
| **P56** | *gdmH* internal rev | AGAATTATCGGAAACGCTC | *gdmH* |
| **P57** | Δ*gdmH* verification for | ATCTCTTGAACATGTGAGCC | slr1142 |
| **P58** | Kol PCR 1080/81 for | CCACTGTTACTGCCAAGAAAC | hypB gene 5'UTR |
| **P59** | Kol PCR 1080/81 rev | CGTCACCCTTTGTCCATCC | nrtD gene |
| **P60** | $P_{sll1080}$ for | AAAGGTCTCGGATGTTTAAAAAGTCAAAATCAACACAGTAAAAC | $P_{sll1080}$ |
| **P61** | $P_{sll1080}$ rev | AAAGGTCTCACGATTTTTCTCCGCTAGTTGTTAATAAC | $P_{sll1080}$ |
| **P62** | $P_{sll1080}$(M) for | AAAGGTCTCGGATGTTTAAAAAATCAAAATCAACAAAGTAAAAC | $P_{sll1080}$ |
| **P63** | sfGFP for 3 | AAAGGTCTCAATCGATGAGCAAAGGAGAAGAACTTTTC | *sfgfp* |
| **P64** | sfGFP rev 2 | AAAGGTCTCGCTTTGGATTTGTCCTACTCAGGAGAGC | *sfgfp* |
| **P65** | p_BB_pUK21_for Pos3.0 | AAACCATGGGAAGACAAAAAGCAGAAGAGACCGGCGGGTGTGGTGGTTA | *pUK21* |
| **P66** | p_BB_pUK21_rev Pos3.0 | AAAACGCGTGAAGACAACATCTTTTGAGACCATTGCGTTGCGCTCACTG | *pUK21* |
| **P67** | EL1-3 for | AAAGAAGACAGAATTGGTCTCAGCCATCGGTCACATGTGCATCCTCGATCTCACTAGTGAGACCGTGTGGTGGTTACGCGCAG | *pUK21* |
| **P68** | EL 4-7 for | AAAGAAGACAGAATTGGTCTCACAGATCGGTCACATGTGCATCCTCGATCTCACATCTGAGACCGTGTGGTGGTTACGCGCAG | *pUK21* |
| **P69** | EL rev | AAAGAAGACATAATTGCGTTGCGCTCACTG | *pUK21* |
| **P70** | pGGC208-CmR LguI for | TTAGCTCTTCTAAATTTGACTTTTGTCCTTTTCCG | pGGC208 |
| **P71** | pGGC208-CmR LguI rev | TTAGCTCTTCGCATTTAGCTTCCTTAGCTCCTG | pGGC208 |
| **P72** | SpecR LguI for | TTAGCTCTTCGATGTCGCGCAGG | pGGC 22 |
| **P73** | SpecR LguI rev | TTAGCTCTTCCTTTCTAGATTTTAATGCGGATG | pGGC 22 |
| **P74** | CEW879 | ATTAATACGACTCACTATAGGATATTGTTTCTAGGGTTCCGATTCCTGGTGATGTGGGGAATGGCTGGTCCGAGAGAAAC | forward oligo for RF00442-WT |
| **P75** | CEW880 | ATTAATACGACTCACTATAGGATATTGTTTCTAGCGTTCCGATTCCTGGTGATGTGGGGAATGGCTGGTCCGAGAGAAAC | forward oligo for RF00442-M1 |
| **P76** | CEW881 | GTCTCCCGGGCTTTTGTCCCGCCGTGTAGCTAGTGGCAAAAATCTTTACAAAGCCTAGCCTGTTTCTCTCGGACCAGCCA | reverse oligo for: RF00442-WT & M1 |

**Table S3: List of plasmids used and generated in this study.**

| Vector | Level | 5' overhang | 3' overhang | Backbone | Selection | Characteristics | Reference |
|---|---|---|---|---|---|---|---|
| **pGGC 0** | 0 | GATG | AAAG | pUC18 | Amp$^R$ | Level 0 empty entry vector based on pUC18 vector with additional integrated *BpiI/BsaI* restriction sites flanking *lacZα* | (9) |
| **pGGC 1** | 1 | GCCA | GTTA | pUK21 | Kan$^R$ | Level 1 position 1 to 6; empty entry vectors based on pUK21 vector with additional integrated BsaI/BpiI restriction sites flanking lacZα | (9) |
| **pGGC 3.0** | 1 | CTAG | CAGA | pUK21 | Kan$^R$ | | This study |
| **pGGC 4** | 1 | CAGA | TGTG | pUK21 | Kan$^R$ | | (9) |
| **pGGC 5** | 1 | TGTG | GAGC | pUK21 | Kan$^R$ | | |
| **pGGC 6** | 1 | GAGC | AGGA | pUK21 | Kan$^R$ | | |
| **pGGC 12** | 0 | GATG | AAAG | pGGC 0 | Amp$^R$ | Level 0 $T_{psbC}$ | |
| **pGGC 22** | 1 | TGTG | GAGC | pGGC 5 | Kan$^R$ | Level 1 position 5 Spec$^R$ | |
| **pGGC 40** | 1 | GCCA | GTTA | pUK21 | Kan$^R$ | End-linker level 1 spanning position 1 till 2 based on pUK21 vector with additional integrated *BsaI/BpiI* restriction sites flanking end-linker sequence TCGGTCACATGTGCATCC TCGATCTCA | |
| **pGGC 42** | 1 | CAGA | CATC | pUK21 | Kan$^R$ | End-linker level 1 spanning position 4 till 7 based on pUK21 vector with additional integrated *BsaI/BpiI* restriction sites flanking end-linker sequence TCGGTCACATGTGCATCC TCGATCTCA | This study |
| **pGGC 46** | 2 | GCCA | CATC | pBluescript II SK (+) | Amp$^R$ | Level 2 empty entry vector based on pBluescript II SK (+) vector with additional integrated *BsaI* restriction sites flanking *lacZα* | (9) |
| **pGGC 47** | 1 | AGGA | CATC | pUK21 | Kan$^R$ | End-linker level 1 spanning position 7 till 7 based on pUK21 vector with additional integrated *BsaI/BpiI* restriction sites flanking end-linker sequence TCGGTCACATGTGCATCC TCGATCTCA | (9) |
| **pGGC 48** | 2 | GCCA | CATC | pBluescript II SK (+) | Cmc$^R$ | Level 2 empty entry vector based on pBluescript II SK (+) vector with additional integrated *BsaI* restriction sites flanking *lacZα* | This study |
| **pGGC 51** | 0 | GATG | AAAG | pGGC 0 | Amp$^R$ | Level 0 Kan$^R$ | This study |
| **pGGC 53** | 1 | TGTG | GAGC | pGGC 5 | Kan$^R$ | Level 1 position 5 Kan$^R$ | This study |
| **pGGC 64** | 0 | GATG | AAAG | pGGC 0 | Amp$^R$ | Level 0 Cmc$^R$ | This study |
| **pGGC 78** | P | GTTA | TGTG | pUK21 | Kan$^R$ | Level P empty vector based on pUK21 vector with additional integrated BsaI/BpiI restriction sites flanking lacZα | This study |
| **pGGC 90** | 1 | TGTG | GAGC | pGGC 5 | Kan$^R$ | Level 1 position 5 Kan$^R$::TtonB | This study |
| **pGGC 132** | 0 | GATG | AAAG | pGGC 0 | Amp$^R$ | Level 0 3’ homologous region for sll 1080 interruption | This study |
| **pGGC 133** | 0 | GATG | AAAG | pGGC 0 | Amp$^R$ | Level 0 5’ homologous region for sll 1080 interruption | This study |
| **pGGC 134** | 0 | GATG | AAAG | pGGC 0 | Amp$^R$ | Level 0 3’ homologous region for sll 1081 interruption | This study |

| | | | | | | | |
|---|---|---|---|---|---|---|---|
| **pGGC 135** | 0 | GATG | AAAG | pGGC 0 | Amp$^{R}$ | Level 0 5' homologous region for sll 1081 interruption | This study |
| **pGGC 136** | 1 | CAGA | TGTG | pGGC 4 | Kan$^{R}$ | Level 1 position 4 3' homologous region for sll 1080 interruption | This study |
| **pGGC 137** | 1 | CAGA | TGTG | pGGC 4 | Kan$^{R}$ | Level 1 position 4 3' homologous region for sll 1081 interruption | This study |
| **pGGC 138** | 1 | GAGC | AGGA | pGGC 6 | Kan$^{R}$ | Level 1 position 6 5' homologous region for sll 1080 interruption | This study |
| **pGGC 139** | 1 | GCCA | CTAG | pUK21 | Kan$^{R}$ | End-linker level 1 spanning position 1 till 3 based on pUC19 vector with additional integrated *BsaI/BpiI* restriction sites flanking end-linker sequence TCGGTCACATGTGCATCCTCGATCTCA | This study |
| **pGGC 199** | 0 | GATG | AAAG | pGGC 0 | Amp$^{R}$ | Level 0 *bvmo* from *Acidovorax* sp. CHX100 | (10) |
| **pGGC 203** | 1 | GAGC | AGGA | pGGC 6 | Kan$^{R}$ | Level 1 position 6 5' homologous region for sll 1081 interruption | This study |
| **pGGC 204** | 2 | - | - | pGGC 48 | Cmc$^{R}$ | Hom. Recomb. template for sll 1080 interruption (pGGC 136 + 138 + 90 + 47 + 139) | This study |
| **pGGC 205** | 2 | - | - | pGGC 48 | Cmc$^{R}$ | Hom. Recomb. template for sll 1081 interruption (pGGC 137 + 203 + 22 + 47 + 139) | This study |
| **pGGC 208** | 2 | GCCA | CATC | pSEVA 351 | Cmc$^{R}$ | Level 2 empty entry vector based on pSEVA 351 vector with additional integrated *lacZα* flanked by *BsaI* restriction sites | (9) |
| **pGGC 288** | 2 | GCCA | CATC | pSEVA | Spec$^{R}$ | Level 2 empty entry vector based on pSEVA 451 vector with additional integrated *lacZα* flanked by *BsaI* restriction sites | This study |
| **pGGC 317** | 2 | GCCA | CATC | pSOMA17 | Gent$^{R}$ | Level 2 empty entry vector based on pSOMA17 vector with additional integrated *lacZα* flanked by *BsaI* restriction sites | This study |
| **pGGC 322** | 0 | GATG | AAAG | pGGC 0 | Amp$^{R}$ | Level 0 P$_{sll1080}$::*sfGFP* | This study |
| **pGGC 323** | 0 | GATG | AAAG | pGGC 0 | Amp$^{R}$ | Level 0 P$_{sll1080}$*(M)*::*sfGFP* | This study |
| **pGGC 324** | 1 | CTAG | CAGA | pUK21 | Kan$^{R}$ | Level 1 Position 3 P$_{sll1080}$::*sfGFP* | This study |
| **pGGC 325** | 1 | CTAG | CAGA | pUK21 | Kan$^{R}$ | Level 1 Position 3 P$_{sll1080}$*(M)*::*sfGFP* | This study |
| **pGGC 326** | 2 | - | - | pSEVA 351 | Cmc$^{R}$ | pSEVA451_ P$_{sll1080}$::*sfGFP* | This study |
| **pGGC 327** | 2 | - | - | pSEVA 351 | Cmc$^{R}$ | pSEVA451_ P$_{sll1080}$*(M)*::*sfGFP* | This study |
| **pGGC 335** | 0 | GATG | AAAG | pUC18 | Amp$^{R}$ | Level 0 Kan$^{R}$::TtonB | This study |
| **pGGC 336** | 1 | AAAG | CAGA | pUK21 | Kan$^{R}$ | Level 1 Position 3.0 TpsbC | This study |
| **pAI 140** | P | GTTA | TGTG | pUK21 | Kan$^{R}$ | *gdmH* deletion template | This study |
| **pAI99** | - | - | - | pSEVA 351 | Cmc$^{R}$ | pSEVA351_PJ$_{23100}$::R$_{Gd}$::*sfgfp*::T*psbC* | This study |
| **pAI100** | - | - | - | pSEVA 351 | Cmc$^{R}$ | pSEVA351_PJ$_{23101}$::R$_{Gd}$::*sfgfp*::T*psbC* | This study |
| **pAI101** | - | - | - | pSEVA 351 | Cmc$^{R}$ | pSEVA351_PJ$_{23109}$::R$_{Gd}$::*sfgfp*::T*psbC* | This study |
| **pAI102** | - | - | - | pSEVA 351 | Cmc$^{R}$ | pSEVA351_PJ$_{23119}$::R$_{Gd}$::*sfgfp*::T*psbC* | This study |
| **pAI103** | - | - | - | pSEVA 351 | Cmc$^{R}$ | pSEVA351_P*cpc*::R$_{Gd}$::*sfgfp*::T*psbC* | This study |
| **pAI104** | - | - | - | pSEVA 351 | Cmc$^{R}$ | pSEVA351_*rhaS*::Prha::R$_{Gd}$::*sfgfp*::T*psbC* | This study |
| **pAI 132** | P | GTTA | TGTG | pUK21 | Kan$^{R}$ | Level P fused up- and downstream region of *gdmH* | This study |
| **pAI225** | - | - | - | pSEVA 351 | Cmc$^{R}$ | pSEVA351_PJ$_{23100}$::R$_{Gd}$::MCS::T*psbC* | This study |

| | | | | | | | |
|---|---|---|---|---|---|---|---|
| **pAI226** | - | - | - | pSEVA 351 | Cmc$^{R}$ | pSEVA351_PJ$_{23101}$::R$_{Gd}$::MCS::T*psbC* | This study |
| **pAI227** | - | - | - | pSEVA 351 | Cmc$^{R}$ | pSEVA351_PJ$_{23109}$::R$_{Gd}$::MCS::T*psbC* | This study |
| **pAI228** | - | - | - | pSEVA 351 | Cmc$^{R}$ | pSEVA351_PJ$_{23119}$::R$_{Gd}$::MCS::T*psbC* | This study |
| **pAI229** | - | - | - | pSEVA 351 | Cmc$^{R}$ | pSEVA351_P*cpc*::R$_{Gd}$::MCS::T*psbC* | This study |
| **pAI230** | - | - | - | pSEVA 351 | Cmc$^{R}$ | pSEVA351_*rhaS*::Prha::R$_{Gd}$::MCS::T*psbC* | This study |
| **pAI231** | - | - | - | pSOMA17 | Gent$^{R}$ | pSOMA17_PJ$_{23100}$::R$_{Gd}$::MCS::T*psbC* | This study |
| **pAI232** | - | - | - | pSOMA17 | Gent$^{R}$ | pSOMA17_PJ$_{23101}$::R$_{Gd}$::MCS::T*psbC* | This study |
| **pAI233** | - | - | - | pSOMA17 | Gent$^{R}$ | pSOMA17_PJ$_{23109}$::R$_{Gd}$::MCS::T*psbC* | This study |
| **pAI234** | - | - | - | pSOMA17 | Gent$^{R}$ | pSOMA17_PJ$_{23119}$::R$_{Gd}$::MCS::T*psbC* | This study |
| **pAI235** | - | - | - | pSOMA17 | Gent$^{R}$ | pSOMA17_P*cpc*::R$_{Gd}$::MCS::T*psbC* | This study |
| **pAI236** | - | - | - | pSOMA17 | Gent$^{R}$ | pSOMA17_*rhaS*::Prha::R$_{Gd}$::MCS::T*psbC* | This study |
| **pAI243** | - | - | - | pSOMA17 | Gent$^{R}$ | pSOMA17_PJ$_{23119}$::R$_{Gd}$::bvmo::T*psbC* | This study |
| **pSEVA351** | - | - | - | pSEVA 351 | Cmc$^{R}$ | Replicative plasmid based on RSF1010 | (11) |
| **pSOMA17** | - | - | - | pSOMA17 | Gent$^{R}$ | Replicative plasmid based on the *Synechocystis* plasmid pCB2.4 | (12) |
| **pSHDY_*P$_{rhaBAD}$*::*mVenus _PJ23119-rhaS*** | - | - | - | pSHDY | Spec$^{R}$ | *P$_{J23119}$*::*rhaS*, *P$_{rhaBAD}$* | (4) |
| **pXG10_SF** | - | - | - | pXG | Cmc$^{R}$ | pSC101-based plasmid containing the *sfgfp* reporter gene | (13) |
| **pSEVA351_PJ23101::RF00442$^{WT}$::*sfgfp*** | - | - | - | pSEVA 351 | Cmc$^{R}$ | PJ23101::RF00442$^{WT}$::*sfgfp* construct integrated into pSEVA351 | This study |
| **pSEVA351_PJ23101::RF00442$^{G16C}$::*sfgfp*** | - | - | - | pSEVA 351 | Cmc$^{R}$ | PJ23101::RF00442$^{G16C}$::*sfgfp* construct integrated into pSEVA351 | This study |

**Table S4: Modular cloning approach for guanidine riboswitch plasmid series.** The plasmids were generated using a BsaI-based GGA in which the given elements were fused via their BsaI cutting sites. Elements inserted in the reactions as plasmids from the GGC collection are given with their GGC number (pGGC xxx). Elements added as PCR products with prior subcloning into pMiniT2.0 are referred to as PCR with the mentioning of the used primer pair as Pxx/Pxx and the used DNA template in brackets. In case of the MCS fragment a PCR without a template yielding primer dimers resembling the MCS with respective BsaI sites was carried out.

| **Plasmid name** | **backbone** | **Position 1** | **Position 2.1 - promoter** | **Position 2.2 – riboswitch** | **Position 2.3 – insert** | **Position 3** | **Position 4-7** |
|---|---|---|---|---|---|---|---|
| pAI99 – pSEVA351_P$J_{23100}$::$R_{Gd}$::*sfgfp*::T*psbC* | pGGC 208 | pGGC 40 | PCR P36/P43; (pSEVA351_ PJ23101::RF00442$^{WT}$::*sfgfp)* | | PCR P44/P45; (pXG10_ SF) | pGGC 336 | pGGC 42 |
| pAI100 – pSEVA351_P$J_{23101}$::$R_{Gd}$::*sfgfp*::T*psbC* | pGGC 208 | pGGC 40 | PCR P37/P43; (pSEVA351_ PJ23101::RF00442$^{WT}$::*sfgfp)* | | PCR P44/P45; (pXG10_ SF) | pGGC 336 | pGGC 42 |
| pAI101 – pSEVA351_P$J_{23109}$::$R_{Gd}$::*sfgfp*::T*psbC* | pGGC 208 | pGGC 40 | PCR P38/P43; (pSEVA351_ PJ23101::RF00442$^{WT}$::*sfgfp)* | | PCR P44/P45; (pXG10_ SF) | pGGC 336 | pGGC 42 |
| pAI102 – pSEVA351_P$J_{23119}$::$R_{Gd}$::*sfgfp*::T*psbC* | pGGC 208 | pGGC 40 | PCR P39/P43; (pSEVA351_ PJ23101::RF00442$^{WT}$::*sfgfp)* | | PCR P44/P45; (pXG10_ SF) | pGGC 336 | pGGC 42 |
| pAI103 – pSEVA351_P*cpc*::$R_{Gd}$::*sfgfp*::T *psbC* | pGGC 208 | pGGC 40 | PCR P34/P35 (Synechocystis gDNA) | PCR P42/P43; (pSEVA351_ PJ23101::RF00442$^{WT}$::*sfgfp)* | PCR P44/P45; (pXG10_ SF) | pGGC 336 | pGGC 42 |
| pAI104 – pSEVA351_*rhaS*::Prha::$R_{Gd}$::*sf gfp*::T*psbC* | pGGC 208 | PCR P46/P47 (pSHDY_Prha$^{BAD}$:: mVenus _PJ23119- rhaS) | PCR P40/P41 (pSHDY_Prha$^{BAD}$:: mVenus _PJ23119- rhaS) | PCR P42/P43; (pSEVA351_ PJ23101::RF00442$^{WT}$::*sfgfp)* | PCR P44/P45; (pXG10_ SF) | pGGC 336 | pGGC 42 |
| pAI225 – pSEVA351_P$J_{23100}$::$R_{Gd}$::MCS::T*psbC* | pGGC 208 | pGGC 40 | PCR P36/P43; (pSEVA351_ PJ23101::RF00442$^{WT}$::*sfgfp)* | | PCR P48/P49 | pGGC 336 | pGGC 42 |
| pAI226 – pSEVA351_P$J_{23101}$::$R_{Gd}$::MCS::T*psbC* | pGGC 208 | pGGC 40 | PCR P37/P43; (pSEVA351_ PJ23101::RF00442$^{WT}$::*sfgfp)* | | PCR P48/P49 | pGGC 336 | pGGC 42 |
| pAI227 – pSEVA351_P$J_{23109}$::$R_{Gd}$::MCS::T*psbC* | pGGC 208 | pGGC 40 | PCR P38/P43; (pSEVA351_ PJ23101::RF00442$^{WT}$::*sfgfp)* | | PCR P48/P49 | pGGC 336 | pGGC 42 |
| pAI228 – pSEVA351_P$J_{23119}$::$R_{Gd}$::MCS::T*psbC* | pGGC 208 | pGGC 40 | PCR P39/P43; (pSEVA351_ PJ23101::RF00442$^{WT}$::*sfgfp)* | | PCR P48/P49 | pGGC 336 | pGGC 42 |
| pAI229 – pSEVA351_P*cpc*::$R_{Gd}$::MCS::T *psbC* | pGGC 208 | pGGC 40 | PCR P34/P35 (Synechocystis gDNA) | PCR P42/P43; (pSEVA351_ PJ23101::RF00442$^{WT}$::*sfgfp)* | PCR P48/P49 | pGGC 336 | pGGC 42 |
| pAI230 – pSEVA351_*rhaS*::Prha::$R_{Gd}$::M CS::T*psbC* | pGGC 208 | PCR P46/P47 (pSHDY_Prha$^{BAD}$:: mVenus _PJ23119- rhaS) | PCR P40/P41 (pSHDY_Prha$^{BAD}$:: mVenus _PJ23119- rhaS) | PCR P42/P43; (pSEVA351_ PJ23101::RF00442$^{WT}$::*sfgfp)* | PCR P48/P49 | pGGC 336 | pGGC 42 |
| pAI231 – pSOMA17_P$J_{23100}$::$R_{Gd}$::MCS::T*psbC* | pGGC 317 | pGGC 40 | PCR P36/P43; (pSEVA351_ PJ23101::RF00442$^{WT}$::*sfgfp)* | | PCR P48/P49 | pGGC 336 | pGGC 42 |
| pAI232 – pSOMA17_P$J_{23101}$::$R_{Gd}$::MCS::T*psbC* | pGGC 317 | pGGC 40 | PCR P37/P43; (pSEVA351_ PJ23101::RF00442$^{WT}$::*sfgfp)* | | PCR P48/P49 | pGGC 336 | pGGC 42 |
| pAI233 – pSOMA17_P$J_{23109}$::$R_{Gd}$::MCS::T*psbC* | pGGC 317 | pGGC 40 | PCR P38/P43; (pSEVA351_ PJ23101::RF00442$^{WT}$::*sfgfp)* | | PCR P48/P49 | pGGC 336 | pGGC 42 |
| pAI234 – pSOMA17_P$J_{23119}$::$R_{Gd}$::MCS::T*psbC* | pGGC 317 | pGGC 40 | PCR P39/P43; (pSEVA351_ PJ23101::RF00442$^{WT}$::*sfgfp)* | | PCR P48/P49 | pGGC 336 | pGGC 42 |
| pAI234 – pSOMA17_P*cpc*::$R_{Gd}$::MCS::T *psbC* | pGGC 317 | pGGC 40 | PCR P34/P35 (Synechocystis gDNA) | PCR P42/P43; (pSEVA351_ PJ23101::RF00442$^{WT}$::*sfgfp)* | PCR P48/P49 | pGGC 336 | pGGC 42 |
| pAI235 – pSOMA17_*rhaS*::Prha::$R_{Gd}$::M CS::T*psbC* | pGGC 317 | PCR P46/P47 (pSHDY_Prha$^{BAD}$:: mVenus _PJ23119- rhaS) | PCR P40/P41 (pSHDY_Prha$^{BAD}$:: mVenus _PJ23119- rhaS) | PCR P42/P43; (pSEVA351_ PJ23101::RF00442$^{WT}$::*sfgfp)* | PCR P48/P49 | pGGC 336 | pGGC 42 |

### Supplementary descriptions

*Expansion of the GGC library*

For BpiI based fusion of PCR products and combination of several genetic cassettes, the level P entry vector pGGC 78 was created analogously to the generation of level 1 plasmids described previously (9) using primer pairs P22/P23 and P24/P25. A kanamycin resistance cassette was integrated into pGGC 0 from a pUK21 plasmid via PCR with primer pair P26/P27 generating pGGC 51 and subsequently transferred into pGGC 5 to produce pGGC 53. The Kanamycin cassette in level 1 (pGGC 53) was expanded by addition of the *tonB* terminator (14) via PCR with primer pair P28/P29 and circulation of the PCR product in a BpiI-mediated Golden Gate assembly yielding pGGC 90. Furthermore, the Kan$^{R}$::T*tonB* construct was re-introduce into pGGC 0 using a PCR with primer pair P7/P8 giving access to pGGC 335. A chloramphenicol resistance cassette was integrated into pGGC 0 via a BsaI-mediated GGA of the PCR product produced with the primer pair P30/P31 from the template pGGC 208 yielding pGGC 64.

The modified pBluescript level 2 entry plasmid with a Cmc resistance (pGGC 48) was produced by a BpiI-based GGA of an amplicon of the pGGC 46 plasmid produced with primer pair P32/P33 omitting the Amp resistance gene and the Cmc$^{R}$ cassette from pGGC 64. A level 2 entry plasmid based on the pSOMA 17 plasmid (Opel et al. 2022) was created by a LguI-mediated GGA of the PCR products produced with primer pair P50/P51 and pSOMA17 as template and primer pair P52/P53 with pGGC 208 as template. To build the level 2 entry vector pGGC 288, the Cmc$^{R}$ cassette of pGGC 208 was replaced with a Spec$^{R}$ cassette using a LguI-mediated GGA of the PCR product generated from pGGC 208 as template with primer pair P70/P71 and the PCR product of primer pair P72/P73 with pGGC 22 as template.

The additional level 1 entry vector pGGC 3.0 and endlinker plasmids pGGC 42 and pGGC 139 were constructed as described for analogous vectors in Lupacchini et al 2025 (9) using the primer pair P65/P66 or P67/P69 and P68/P69, respectively. The newly created pGGC 3.0 was subsequently used to integrate the terminator of psbC from pGGC 12 into level 1 position 3.0 generating pGGC 336.

*Cloning of the plasmid series for guanidine-dependent gene expression*

For the creation of the plasmid series featuring the combination of different promoters with the guanidine riboswitch and either *sfgfp* or a multi cloning site (MCS), a modular Golden Gate approach using plasmids from the GGC collection and PCR products integrated into pMiniT2.0

was applied. With that approach, the plasmids were constructed analogously to a level 2 assembly. Thereby, position 1 was filled with either an endlinker sequence from pGGC 40 or in case of plasmids with the rhamnose-inducible promoter with a PJ23119::*rhaS* cassette. At position 2 the respective combinations of a promoter, the RF00442 sequence, and either *sfgfp* or an MCS with BsaI-cutting sites for scarless joining were inserted. The transcriptional terminator TpsbC from pGGC 336 was appointed to position 3 and the endlinker sequence from pGGC 42 was used to span from position 4 to the backbone. As level 2 entry vectors, pGGC 208 was used for the pSEVA351-based plasmids and pGGC 317 for the pSOMA17-based plasmids.

*Protein extraction and immunoblotting*

Samples of *Synechocystis* cells were taken 24h after the addition of 1.25 mM guanidine-HCl and concentrated to an $OD_{750}$ of 4.5 in a volume of 1 ml. Protein extraction, SDS-PGAE and Western blot were performed as described previously (65) with the following exceptions: The protein concentration in the extracts was estimated by measuring their absorbance at 280 nm and 260 nm using the formula c[µg/µl] = 1.55 x $A_{280}$ – 0.76 x $A_{260}$; Based on that, 7.5 µg of protein were loaded per lane onto the SDS-PAGE; An Anti-His-HRP conjugate antibody (Qiagen) was used for immunodetection; Lumi-Light$^{PLUS}$ (Roche) Western-Blot-Substrate was used for chemiluminescence detection performed in a FUSION FX7 EDGE imaging system.

*Berkley Madonna script used to simulate a guanidine-dependent steady-state bioprocess*

```
METHOD RK4

STARTTIME = 0
STOPTIME=145
DT = 0.1

{Balances}
V' = D - O                          ; Volume balance
X'= X*U - O*cX                      ; Biomass balance
VD' = D                             ; Feed
VO'=O                               ; Outflow
S' = D*cfS - R - O*cS               ; Substrate balance
P' = R - O*cP                       ; Product balance
N' = X*U*-YXN*Gp - O*cN             ; Nitrate balance
G' = D*cfG - X*U*YXG - O*cG         ; Guanidine balance
PO' = O*cP                          ; Product outflow
SO' = O * cS                        ; Substrate outflow

{Initial Conditions}
INIT X=X0
INIT V=V0
INIT VD = 0
INIT VO = 0
INIT N = N0
INIT G = G0
INIT S = 0
INIT P = 0
INIT PO = 0
Init SO = 0
INIT Ag = 0
```

```
{amounts to concentrations}
cN = N/V
cX=X/V
cS = S/V
cP = P/V
cG = G/V
cGm = cG/59.07

{Kinetics}

R = If S>0 then R1 else 0
R1 = A*X
D= If X>Xt then F else 0
O= If V>V0 then F else 0
Gp = If cG<=0 then 1 else 0

{Enzyme activity}
A = Aconst + Ag
Ag' = ((Aind * cGm)/(0.000001 + cGm))/24          ; Addition of enzyme activity over 24 after Guanidine addition based on a
                                                   Michaelis-Menten like kinetic
Limit Ag <= Aind

{limits}
LIMIT N >=0
LIMIT G >=0
LIMIT S >=0


{Constants}
U=0.05                          ; Maximal specific growth rate, 1/h
X0=0.1/4.5*V0                   ; Inoculum biomass (g)
V0=1                            ; Start volume
Xt=1/4.5*V0                     ; Targeted biomass (g)
F=1                             ; Pump rate (l/h)
Aind = 3.8              ; Enzyme activity inducible (U/gcdw = mmol/h*gCDW)
Aconst = 0.4                    ; Enzyme activity uninduced
cfS = 15                ; Concentration of Substrate in Feed (mM)
cfG = 0.295                     ; Concentration of Guanidine in Feed (g/l)
N0 = 0.017*62*V0        ; Nitrate start amount (g)
YXN = 0.8               ; Yield biomass on nitrate (g nitrate for g biomass)
G0 = 0                          ; Start amount of Guanidine (g)
YXG = 0.256                     ; Yield biomass on guanidine (g guanidine for g biomass)

{variables}
;cX biomass concentration (g/l)
;X biomass amount (g)
;D dilution rate (l/h)
;O outflow rate (l/h)
;V volume (l)
;F feed rate (l/h)
;VD total volume for dilution
;S substrate amount (mmol)
;cS substrate concentration (mM)
;P product amount (mmol)
;cP product concentration (mM)
;A enzyme activity
;Ag guanidine based enzyme activity
;cG Guanidine concentration (M)
;cN nitrate concentration (mM)
;Gp Guanidine priorization --> if guanidine is present no nitrate is used
```

### Supplementary References


1. S. C. Albers, V. A. Gallegos, C. A. M. Peebles, Engineering of genetic control tools in *Synechocystis* sp. PCC 6803 using rational design techniques. *J. Biotechnol.* **216**, 36–46 (2015).

2. H.-H. Huang, P. Lindblad, Wide-dynamic-range promoters engineered for cyanobacteria. *J. Biol. Eng.* **7**, 10 (2013).

3. A. J. Victoria, *et al.*, A toolbox to engineer the highly productive cyanobacterium *Synechococcus* sp. PCC 11901. *Plant Physiol.* **196**, 1674–1690 (2024).

4. A. Behle, P. Saake, A. T. Germann, D. Dienst, I. M. Axmann, Comparative dose-response analysis of inducible promoters in cyanobacteria. *ACS Synth. Biol.* **9**, 843–855 (2020).

5. C. M. Immethun, *et al.*, Physical, chemical, and metabolic state sensors expand the synthetic biology toolbox for *Synechocystis* sp. PCC 6803. *Biotechnol. Bioeng.* **114**, 1561–1569 (2017).

6. B. Blasi, L. Peca, I. Vass, P. B. Kós, Characterization of stress responses of heavy metal and metalloid inducible promoters in *Synechocystis* PCC6803. *J. Microbiol. Biotechnol.* **22**, 166–169 (2012).

7. L. M. Briggs, V. L. Pecoraro, L. McIntosh, Copper-induced expression, cloning, and regulatory studies of the plastocyanin gene from the cyanobacterium *Synechocystis* sp. PCC 6803. *Plant Mol. Biol.* **15**, 633–642 (1990).

8. Y. Nakahira, A. Ogawa, H. Asano, T. Oyama, Y. Tozawa, Theophylline-dependent riboswitch as a novel genetic tool for strict regulation of protein expression in cyanobacterium *Synechococcus elongatus* PCC 7942. *Plant Cell Physiol.* **54**, 1724–1735 (2013).

9. S. Lupacchini, *et al.*, Co-expression of auxiliary genes enhances the activity of a heterologous $O_2$-tolerant hydrogenase in the cyanobacterium *Synechocystis* sp. PCC 6803. *Biotechnol. Biofuels Bioprod.* **accepted** (2025).

10. A. Tüllinghoff, H. Djaya-Mbissam, J. Toepel, B. Bühler, Light-driven redox biocatalysis on gram-scale in *Synechocystis* sp. PCC 6803 via an *in vivo* cascade. *Plant Biotechnol. J.* **21**, 2074–2083 (2023).

11. R. Silva-Rocha, *et al.*, The Standard European Vector Architecture (SEVA): a coherent platform for the analysis and deployment of complex prokaryotic phenotypes. *Nucleic Acids Res.* **41**, D666-675 (2013).

12. F. Opel, *et al.*, Generation of synthetic shuttle vectors enabling modular genetic engineering of cyanobacteria. *ACS Synth. Biol.* (2022). https://doi.org/10.1021/acssynbio.1c00605.

13. C. P. Corcoran, *et al.*, Superfolder GFP reporters validate diverse new mRNA targets of the classic porin regulator, MicF RNA. *Mol. Microbiol.* **84**, 428–445 (2012).

14. K. Postle, R. F. Good, A bidirectional rho-independent transcription terminator between the *E. coli tonB* gene and an opposing gene. *Cell* **41**, 577–585 (1985).
